# DropBlot: single-cell western blotting of chemically fixed cancer cells

**DOI:** 10.1101/2023.09.04.556277

**Authors:** Yang Liu, Amy E. Herr

**Affiliations:** Department of Bioengineering, University of California, Berkeley, California 94720, USA; Chan Zuckerberg Biohub, San Francisco, California 94158

## Abstract

To further realize proteomics of archived tissues for translational research, we introduce a hybrid microfluidic platform for high-specificity, high-sensitivity protein detection from individual chemically fixed cells. To streamline processing-to-analysis workflows and minimize signal loss, DropBlot serially integrates sample preparation using droplet-based antigen retrieval from single fixed cells with unified analysis-on-a-chip comprising microwell-based antigen extraction followed by chip-based single-cell western blotting. A water-in-oil droplet formulation proves robust to the harsh chemical (SDS, 6M urea) and thermal conditions (98°C, 1-2 hr.) required for sufficient antigen retrieval, and the electromechanical conditions required for electrotransfer of retrieved antigen from microwell-encapsulated droplets to single-cell electrophoresis. Protein-target retrieval was demonstrated for unfixed, paraformaldehyde-(PFA), and methanol-fixed cells. We observed higher protein electrophoresis separation resolution from PFA-fixed cells with sufficient immunoreactivity confirmed for key targets (HER2, GAPDH, EpCAM, Vimentin) from both fixation chemistries. Multiple forms of EpCAM and Vimentin were detected, a hallmark strength of western-blot analysis. DropBlot of PFA-fixed human-derived breast tumor specimens (n = 5) showed antigen retrieval from cells archived frozen for 6 yrs. DropBlot could provide a precision integrated workflow for single-cell resolution protein-biomarker mining of precious biospecimen repositories.

## Introduction

An estimated 1 billion archived tumor tissues are housed in biorepositories and medical centers^1^. Archived tissues make retrospective studies possible and retrospective studies are needed to understand disease and develop therapies. For example, retrospective studies of tissues powered the development of trastuzumab (Herceptin®), arguably one of the most effective targeted cancer therapies ever developed^2^. To preserve cellular morphology and prevent the degradation of proteins during storage, archived tissues are chemically fixed (i.e., formalin^3^; paraformaldehyde (PFA)^4^; methanol^5^; formalin-fixed, paraffin-embedded (FFPE)^6^). Cell fixation is a crucial component of clinical pathology and biomedical research, allowing stabilization of proteins during extended archiving of the cellular material. Prior to analysis of archived cells and tissue, these fixed cells require harsh pretreatments to partially restore antigen immunoreactivity for subsequent measurement (i.e., antigen retrieval)^7–9^ typically by immunoassay. Although much remains to be learned about the design of high-quality tissue-fixation protocols and the mechanisms by which antigen immunoreactivity is restored, fixed and archived biospecimens have substantially influenced clinical medicine, and will continue to do^10, 11^.

For analysis of fixed tissues and even single fixed cells, immunohistochemistry (IHC)^12^ and immunocytochemistry (ICC)^13^ are widely used in pathology and biomedical research labs. Fluorescence and colorimetric stains report protein-target presence, localization, and distribution (e.g., human epidermal growth factor receptor 2 (HER2)^14^, programmed death ligand 1 (PD-L1)^15^). For analysis of suspensions of fixed cells, flow cytometry is a potent tool for rapid protein analysis and cell sorting (i.e., size, shape, and biomolecular profile using fluorescently labeled antibody probes)^16, 17^. Flow cytometry can process more than 10,000 cells per second with multiplexity of up to ∼20 protein targets^16^. While useful, flow cytometry requires a large starting number of cells (>10,000 cells), and limited detection specificity due to inadequate probes, probe and spectral signal overlap, and cellular autofluorescence^16^. In contrast, mass spectrometry (MS) is a powerful protein detection tool that does not require antibody probes, but MS has limitations and is not appropriate for all protein-based questions^18^. Top-down proteomics struggles with single-cell detection^18^, and bottom-up MS obscures proteoform stoichiometry by requiring protein digests as input^19^. Antibody-based MS methods offer high-resolution analyses but inherently depend on specific antibody probes^20^, similar to immunoassays. Proteoform Imaging Mass Spectrometry (PiMS) provides increased spatial resolution but cannot analyze proteins >70 kDa, limiting the use of PiMS to certain cancer-related protein targets^21^.

Prior to any analytical stage, sample preparation is a critical stage with microfluidic techniques making inroads into the preparation of fixed cells. Droplet-based^22^ and reaction chamber-based^23^ microfluidic systems offer an enclosed compartment to facilitate cell incubation, lysis, and antigen extraction under even harsh conditions, in preparation for analysis of diverse cell types and myriad fixation conditions. Microfluidic large-scale integration (mLSI) platforms^24^, single-cell barcode chips (SCBCs)^25^, and droplet-based cell screening & sorting^26^ reduce the number of cells required for protein analysis, as compared to flow cytometry and conventional MS. New measurement methods^27^ may improve protein detection sensitivity and specificity in fixed cell and tissue samples.

Recently adapted to single-cell resolution, a workhorse targeted proteomic method is slab-gel immunoblotting. A type of immunoblot, western blotting couples protein polyacrylamide gel electrophoresis (PAGE) with subsequent immunoassays^28^. Slab-gel immunoblotting of fixed samples has been reported^29, 30^, often pooling tissue or cell samples to enhance detection sensitivity. For fresh or fresh-frozen clinical specimens, our research group has introduced single-cell immunoblotting^31, 32^. To foster technology translation and support concurrent analyses of 100’s-1000’s of cells, we design ‘open-microfluidic’ chips that do not incorporate enclosed microchannels or pneumatic control (pumps, valves). Instead, the open-fluidic chips are similar to a mini-gel layered on a standard microscope slide, but with arrays of open microwells stippled into the open-faced gel. The microwells are sized to isolate single dissociated tumor cells and circulating tumor cells for subsequent imaging, lysis, and ultra-rapid protein PAGE through the polyacrylamide gel surrounding each microwell. While relevant to readily lysed fresh cells, the open-microwell design supports only brief cell-lysis durations (<1 min, <50°C) before the lysate dilutes and diffuses out of the microwell, thus making the readily translatable tools irrelevant to preparation of fixed cells that require long-duration (>60 min) and harsh antigen retrieval conditions^33, 34^. Consequently, we introduce a hybrid microfluidic tool – called DropBlot – that extends the relevance of open-fluidic chip design to the preparation and analysis of single fixed cancer cells.

DropBlot combines droplet-based, single-cell sample preparation with open-fluidic, single-cell western blotting for 100’s of chemically fixed cells. To restore antigen immunogenicity prior to electrophoresis and immunoassays, we report the design of stable water-in-oil (W/O) droplets to encapsulate single fixed cells and support cell lysis at high temperatures (100°C) and long incubation periods (1-2 hrs.). Lysate-containing droplets are sedimented into open microwells. We employ electrotransfer to inject solubilized protein targets from each microwell-encapsulated droplet into the PAGE chip, for protein separation, in-gel protein blotting (covalent immobilization of protein to gel polymer), and subsequent immunoassays. We develop the DropBlot workflow using two breast cancer cell lines and a suite of chemical fixation conditions, and then apply DropBlot to investigate a pilot cancer-protein panel composed of an epithelial marker (EpCAM), a mesenchymal marker (vimentin, VIM), and human epidermal growth receptor 2 (HER2) in a pilot group of breast cancer patient-derived cell samples. Our results demonstrate western blotting of single PFA-fixed cells, and form the basis of a modular sample-preparation and analysis tool, adaptable to the panoply of cell-fixation conditions used in biorepositories.

## Results and Discussion

### Overview of DropBlot

To perform single-cell western blotting on antigen targets retrieved from chemically fixed cells, we sought to seamlessly integrate two disparate functions into one assay workflow: (1) antigen retrieval from single fixed cells using droplet microfluidic technology with (2) single-cell PAGE and immunoblotting of the retrieved antigen targets using a planar array of single-cell PAGE separations on an open microfluidic device (**Figure 1a**). The hybrid DropBlot design is comprised of three typically independent microfluidic modes (i.e., cell-laden droplets, microwells, and planar chip separations) as a way to unify the diverse chemical, thermal, electrical, and mechanical conditions required at each stage of the preparation and analysis process into one integrated workflow (**Figure 1b-f**).

**Figure 1.**
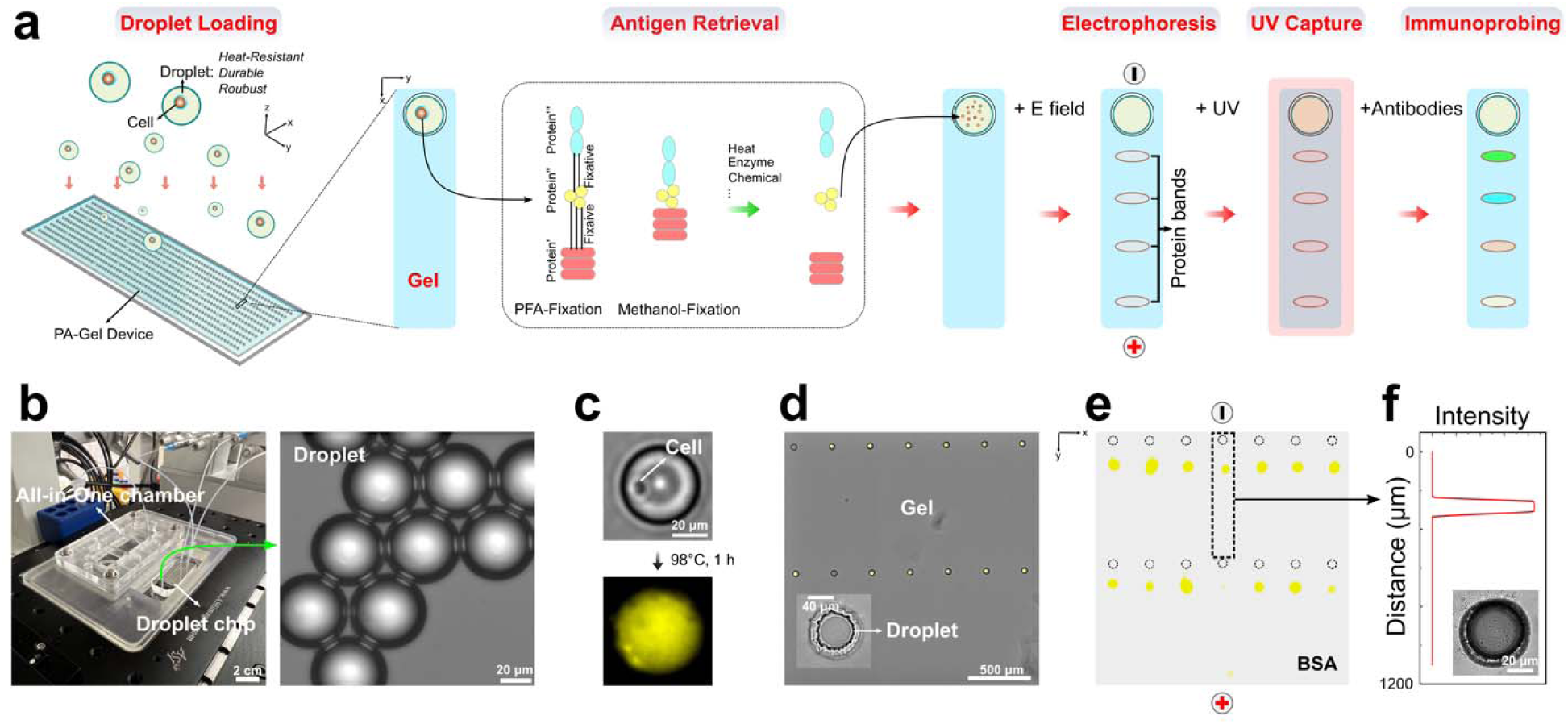
Design, operation, and characterization of DropBlot, a hybrid microfluidic platform that couples single fixed-cell sample preparation in droplets to single-cell protein immunoblotting in a planar, open-fluidic chip. (a) Conceptual schematic of the droplet-based single-cell preparation stage interfacing with the device-based single-cell western blotting stage. The tandem assay is designed to solubilize immunoreactive antigen targets from individual chemically fixed cells for subsequent single-cell immunoblotting. (b) Photo of the hybrid-assay assembly, which mates the droplet-generation chip with a PAGE chamber (left). Brightfield image of droplets for cell preparation stage (right). (c) Fluorescence micrographs of GFP-expressing MCF7 lysing in a W/O droplet. False-color yellow is GFP fluorescence signal. (d) Brightfield micrograph microwell array and abutting PAGE regions for single-cell western blotting. False-color yellow is AF488-labeled BSA signal. Inset: micrograph of a droplet-containing microwell. (e) Fluorescence micrograph of an array of PAGE endpoint analyses of AF488-labeled BSA protein standard (Δt_PAGE_ = 20 s, *E* = 40 V/cm). Anode and cathode orientation is as marked. (f) Background-subtracted fluorescence intensity (AFU) of one PAGE lane. Inset: brightfield micrograph of a droplet-containing microwell after PAGE.

### Design of droplets for stability to the chemical, thermal, and mechanical requirements of DropBlot

We sought to achieve four performance goals by selecting a water-in-oil (W/O) chemistry for the droplets and a microfluidic H-junction for droplet generation. First, we sought to co-encapsulate single cells and a surfactant-containing antigen-retrieval buffer, in such a way that antigen solubilization would occur after the droplet is fully formed. The performance goal seeks to ensure sufficient antigen retrieval while minimizing antigen leakage from the droplet. Second, we sought a droplet chemistry that maintains stability under harsh antigen-retrieval conditions which include the presence of surfactants (SDS), exposure to elevated temperature (>95°C), and incubation for longer than 1 hr (see **Methods**)^35, 36^. Third, we sought a stable-droplet formulation robust to mechanical handling and seating of said droplets into the open-fluidic array of microwells. Fourth, we sought a stable-droplet formulation amenable to electrotransfer of solubilized antigen out of each microwell-encapsulated droplet and into a PAGE gel proximal to each microwell.

Given these four target-performance specifications, we opted for a water-in-oil (W/O) droplet chemistry, with droplets generated at an H-junction to allow co-introduction of single cells with the antigen-retrieval buffer. Our first step was systematically optimizing the droplet generation function for single cell per droplet occupancy, by considering channel geometry, flow rates of the continuous and dispersed phases, and the initial concentration of the cell suspension (MCF7 cells, **Figure S1**). Single-cell occupancy was achieved in 79.5 ± 6.1% (*n* = 3) of droplets generated under the following empirically determined conditions: droplet diameter = Ø*_droplet_* = 40-60 µm; volumetric flow rate of dispersed phase = Q*_dispersed_* = 0.2-20 µL/min; volumetric flow rate of continuous phase = Q*_continuous_* = 0.2-100 µL/min; starting concentration of cell suspension = 1.0-6.0×10^6^ cells/mL.

After establishing baseline single cell per droplet occupancy with the H-junction, we sought to establish stable droplets compatible with our selected antigen-retrieval protocols. For the droplet design rules, we considered two major factors: (i) properties of the surfactants used in the continuous phase (mineral oil)^37, 38^ and (ii) properties of the dispersed phase (dual lysis and antigen-retrieval buffer which includes SDS, cell suspension)^39, 40^. In the continuous phase, surfactants can reduce interfacial tension and prevent droplets from coalescing^37^. For example, mineral oil supplemented with 0.5-5.0% (v/v) of Span® 80 nonionic surfactant results in stable W/O emulsions^41, 42^. The dispersed phase – especially when containing SDS or other surfactants – also influences droplet stability^43, 44^. As is shown in **Figure 2a**, a dispersed phase containing increasing SDS concentration decreases droplet stability (i.e., *Q_dispersed_,=* 10 µL/min, *Q_continuous_*= 15 µL/min) as expected, due to destabilization of hydrophilic surfactants in the dispersed phase^40^. To determine a suitable formulation for our DropBlot performance goals, we screened a panel of formulations for both the continuous (Span® 80 concentrations = 0.5-5% (v/v)) and dispersed phases (SDS concentrations = 0.1-2% (w/v)). We scrutinized the bulk stability of the droplet formulations over a 3-hr incubation at elevated temperature (80-100°C) by employing brightfield imaging of a homogeneous suspension of droplets in a tube (**Figure 2b-c**). Any visually detectable phase separation of the suspension (with the concomitant visible formation of immiscible layers) indicates droplet breakage. We observed no observable phase separation of materials layers for the formulation consisting of a continuous phase containing 2% (v/v) Span® 80 and a dispersed phase containing 0.5% SDS (w/v) (Ø*_droplet_*= 50µm). Further, we observed no notable change in the number of droplets by droplet enumeration microscopy (Δt = 0 hr, *n*_droplets_ = 657 ± 31; Δt = 3 hr, *n*_droplets_ = 588 ± 28; n = 3). Given these findings, we next sought to understand the cell-lysis efficacy for model cell lines. Model cell lines are important for iterative assay development, because primary cells can be precious and highly variable. When engineering to express fluorescent reporter proteins, model cell lines also allow imaging-based assay optimization. Using fluorescence microscopy inspection, we observed cell lysis of GFP-expressing MCF7 cells within 5 min with a 0.5% SDS (w/v) formulation in the dispersed phase (**Figure 2d-e**; 95.1 ± 1.2% lysis efficiency; *n*_droplets_ = 1000; n = 3). Important to our mechanical robustness performance goal, brightfield visual inspection showed lysate-filled droplets remained largely intact after deposition into the microwells (92.3 ± 4.7% yield of intact droplets; *n*_microwells_ = 560; n = 3).

**Figure 2.**
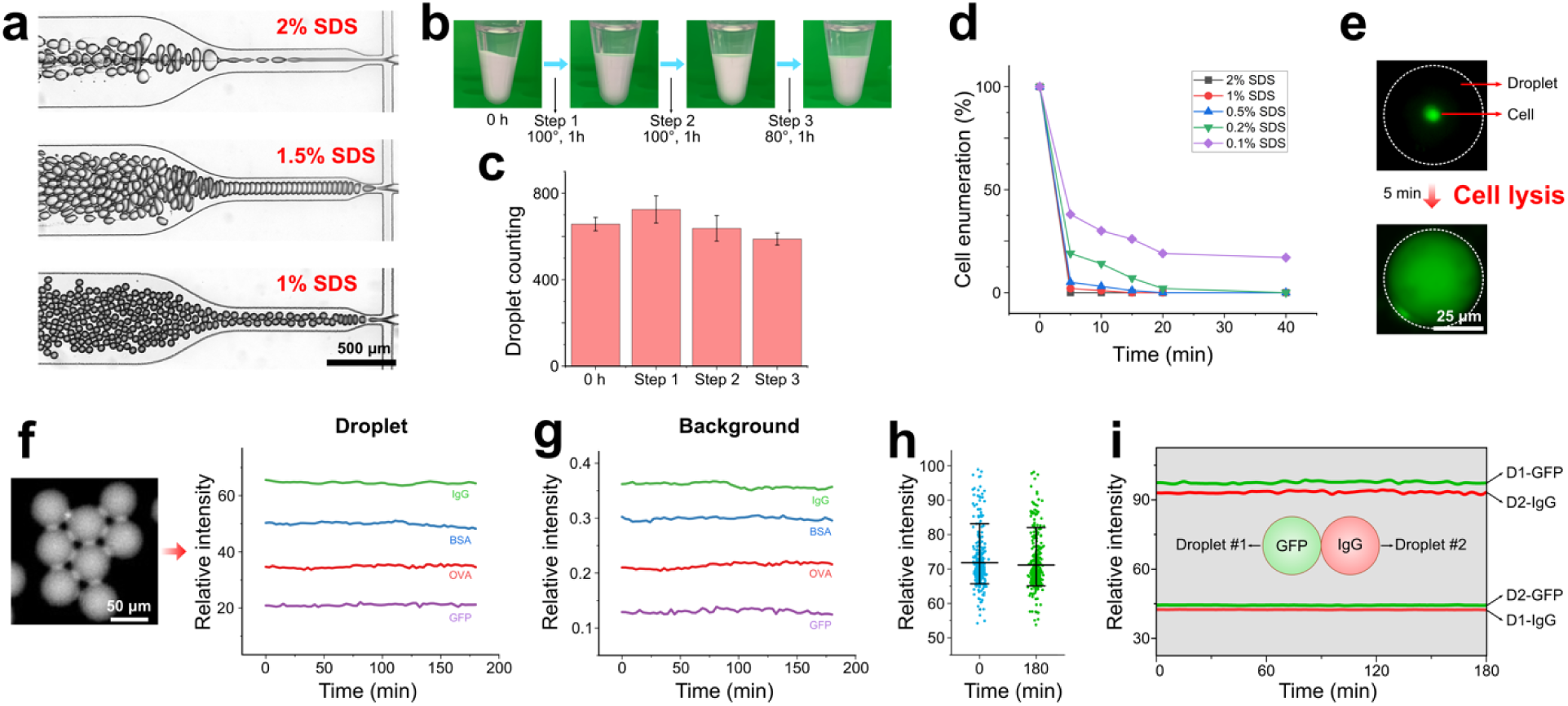
Design and generation of stable droplets as required for fixed-cell lysis and antigen retrieval, followed by droplet loading into individual microwells for electrotransfer to lysate PAGE. (a) Brightfield micrographs show droplet generation with 0.5–2.0% (w/v) SDS in the dispersed medium (V_dispersed_: V_continuous_ = 10: 15 µL/min). Continuous phase [Span80] = 2% (v/v). (b) Brightfield image time series assessing droplet stability for the incubation sequence: Step 1 (100°C for 1 h) ➔ Step 2 (100°C for 1 h) ➔ Step 3 (80°C for 1 h). Top layer (transparent): mineral oil; Bottom layer (opaque): droplets. (Ø_droplet_ = 50 µm, 0.5% (w/v) SDS) (c) Droplet enumeration during the incubation sequence in (b). Droplets were incubated and imaged on a glass slide with a hydrophobic surface. (d) Fraction of MCF7 cells lysed as a function of SDS concentration at 95°C. (Ø_droplet_ = 50 µm) (e) Fluorescence micrograph showing ‘before’ and ‘after’ lysis of a GFP-expressing MCF7 cell encapsulated in a W/O droplet (RT, 0.5% SDS (w/v)). (f) Fluorescence micrograph of a droplet loaded with Alexa-Fluor 555 labeled BSA. (right): Mean fluorescence intensity monitoring of ∼300 individual droplets, each loaded with AF488-IgG, AF555-BSA, AF647-OVA, and GFP. (g) Mean fluorescence intensity monitoring of background signal for the droplets in (f). (h) Mean fluorescence intensity monitoring of 300 single droplets loaded with AF555-BSA. (i) Mean fluorescence intensity of paired droplets loaded with GFP (left droplet, green shading) and AF555-IgG (right droplet, red shading). Flux of fluorescence signal is considered into and out of both droplets. GFP intensity in Droplet #1 (D1-GFP); IgG intensity in Droplet #2 (D2-IgG); GFP intensity in Droplet #2 (D2-GFP); IgG intensity in Droplet #1 (D1-IgG).

To ensure droplet stability under the chemical conditions required by the antigen-retrieval function, we next evaluated protein leakage from the selected W/O droplet formulation using a continuous phase of 2% (v/v) Span® 80 and a dispersed phase containing 0.5% SDS (w/v). A fluorescently labeled protein standard solubilized in antigen-retrieval buffer (SDS: 0.5% (w/v)) was loaded into the droplets, with fluorescence microscopy used to monitor the total integrated fluorescence intensity of individual droplets over time. We observed no notable decrease in fluorescence intensity during the 180-min monitoring period at room temperature (**Figure 2f**). The background intensity remained nearly constant over 3 hrs (**Figure 2g**), and the intensity distribution of individual droplets did not notably differ before and after the experiment (**Figure 2h**). Additionally, we observed no notable fluorescence crosstalk between neighboring droplets loaded with protein ladder targets labeled with spectrally distinct fluorophores (**Figure 2i**). These findings suggest a formulation for stable droplets suitable for the preservation of target proteins with negligible target loss or crosstalk over the course of a 180-min experiment. Taken together, we adopted the described conditions as a starting point for developing the DropBlot for fixed cancer cells.

### Design of droplets for stability to the electrical and mechanical requirements of DropBlot

We next sought to determine suitable conditions for (i) mechanical deposition of the cell-lysate-containing droplets onto the microwell array and (ii) electrotransfer of soluble protein targets from microwell-encapsulated droplets into the proximal polyacrylamide gel for single-cell western blotting of retrieved antigens. To understand the impact of physical handling of droplets, W/O droplets containing lysate from single unfixed cells were mechanically deposited onto the open-fluidic chip used for the DropBlot protein analysis stage. Droplets were sedimented onto the surface of the open gel, and gentle washing removed droplets that did not sediment into microwells (**Figure S2**). Inspection of the microwells by fluorescence microscopy reported ∼95.1 ± 3.1% (n = 3 chips) of microwells occupied with a single droplet after 15 min of settling. We observed that intact droplets loaded into the microwells did not noticeably coalesce with the polyacrylamide gel walls of the microwells.

We next aimed to understand both the feasibility and efficiency of electrotransfer as a low-dispersion means to introduce solubilized (retrieved) antigens from the microwell-encapsulated droplet into the microwell-abutting protein PAGE analysis gel. While the electrotransfer concept is based on an approach our group has used for electrophoretic analysis of single-cell lysates^31^, DropBlot presents a notable difference in the presence of an immiscible phase between the starting location of the antigen targets, that being in a droplet, and the endpoint location, that being in a proximal molecular sieving hydrogel.

To understand this more chemically and geometrically complex sample-injection configuration, we asked how the presence of an immiscible phase and the placement of each droplet within a microwell affects injection of protein target into the sieving matrix, with particular interest in injection efficiency and dispersion (**Figure 3a**, b).

**Figure 3.**
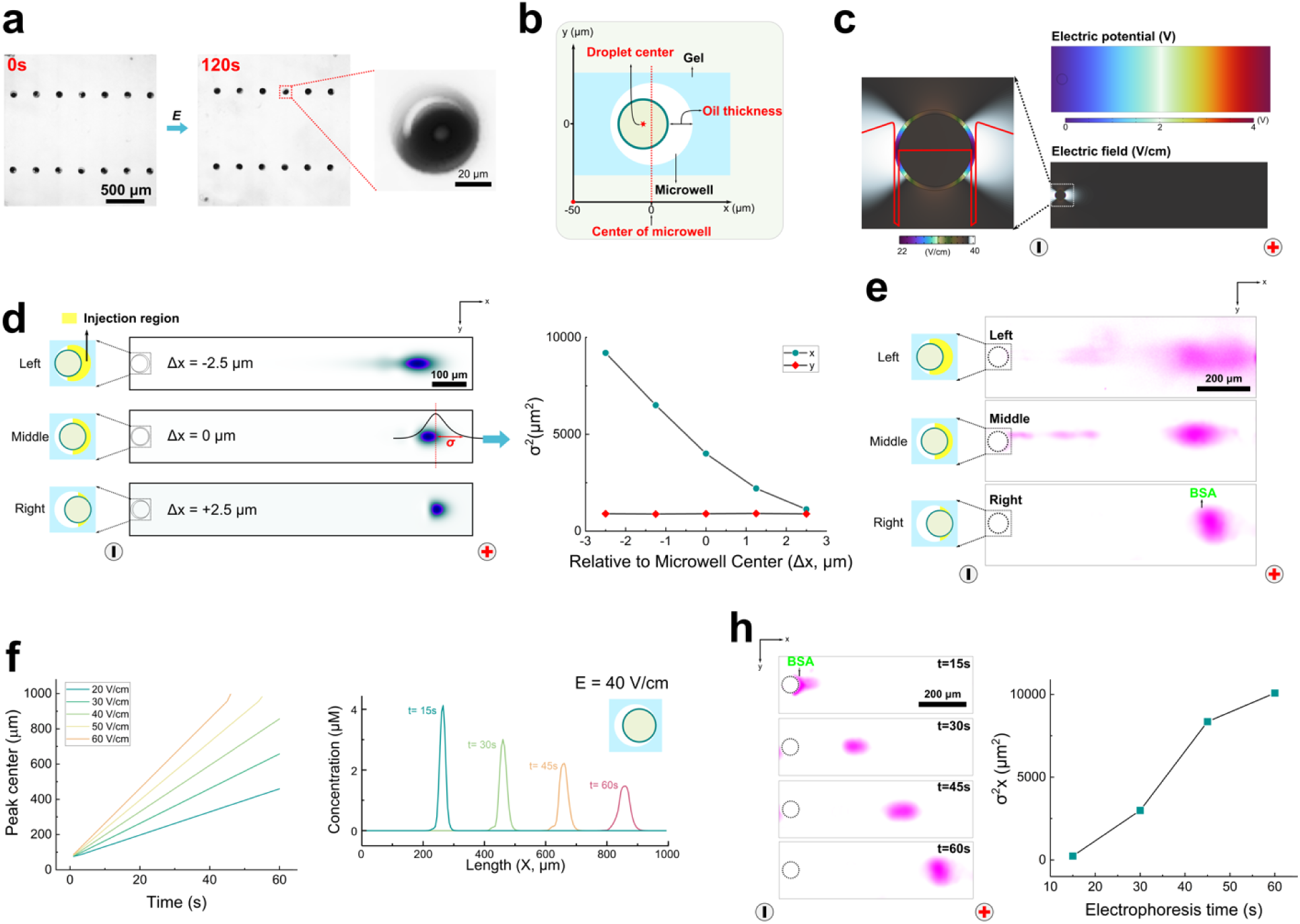
Simulation and validation of protein-target electrotransfer from a microwell-encapsulated W/O droplet into abutting PAGE gel. (a) Droplet stability test under electric field (field strength: 40 V/cm). The droplets remain intact after 120s’ electrophoresis. (b) Schematic showing a top view (x-y) of the planar DropBlot model for study of electrotransfer of protein lysate from droplet, through oil region, into thin-layer polyacrylamide gel for PAGE. Microwell center is located at *x* = 0 µm. Droplet *y* position is aligned with *y*-position of the microwell. (c) Simulation showing the applied potential and resultant electric field around geometry shown in (b). The red line represents electric field strength along the *x*-direction of the PAGE separation gel abutting each microwell. (d) Simulation of the migration distance (left) and concentration profiles (right) of protein ladder component BSA as a function of various droplet positions relative to microwell center (Left: Δx = –2.5 µm; Middle: Δx = 0 µm; Right: Δx = +2.5 µm). Yellow shading demarcates injection region; σ = BSA peak width; Ø_microwell_ = 50 µm; Ø_droplet_ = 45 µm; E = 40 V/cm; Δt_PAGE_ = 60 s. (e) Companion fluorescence micrographs of protein ladder component AF555-BSA for the same conditions presented in simulations reported in (d). (f) Simulation of Right-aligned droplet (Δx = +2.5 µm) configuration and BSA PAGE electromigration as a function of elapsed PAGE time (*E* = 40 V/cm) and applied electric field strength. (h) Companion fluorescence micrographs of AF555-BSA electromigration along the PAGE separation axis for a Right-aligned droplet (Δx = +2.5 µm) configuration. Resultant empirical relationship between concentration profile and electrophoresis time.

Using a 2D numerical simulation that we created in COMSOL®, we considered the immiscible and miscible phases (i.e., oil layer thickness) and geometry (i.e., electric field strength, microwell size, relative position between droplet and microwell). The conductivity of the oil layer was assumed to be similar to mineral oil, σ = 0.175 S/m (note: the conductivity of Span® 80 is negligible compared to mineral oil^45, 46^). Simulations suggest that a voltage drop across the droplet results in an applied electric field within the core of the droplet and through the immiscible oil layer (**Figure 3c**). Establishing a continuous – if not uniform – electric field across all phases of the configuration is the minimum necessary attribute needed to use electrotransfer to move a charged analyte from within a droplet, through the droplet, and into an abutting polyacrylamide gel.

The 2D simulations further informed us that the applied electric field distribution across the droplet-to-microwell interface would create an analyte-stacking interface (at the O/W interface), and an analyte-destacking interface (as the species electromigrate through the oil layer). A second analyte-staking interface would be expected upon analyte electromigration into the polyacrylamide gel interface forming the walls of each microwell. To understand and control these discrete regions, we define an ‘injection region’ as the span between the right edge of a microwell-encapsulated droplet and the right edge of the encapsulating microwell. When the anode is located to the right along the separation axis, the protein-migration direction is defined as +*x* (i.e., negatively charged protein electromigrates from left to right, **Figure 3d**). For a fixed microwell diameter, the thickness of the injection region is determined by the droplet diameter and the relative position of the droplet in the microwell (Δx). The retention time in the injection region, is estimated by *t*_RT_ = *L*_inj_ / (*E*_inj_ × μ_inj_) where *L*_inj_ is the length of the injection region (cm), *E*_inj_ is the strength of the applied electric field (V/cm) in the injection region, and the μ_inj_ i the electrophoretic mobility (cm^2^/(V·s)) of the charged analyte through the injection region (e.g., =125 µm^2^/(V·s); **Figure S3**)

Given the nonuniform electric field distribution across the droplet-to-microwell geometry – and anticipated concomitant impact on peak dispersion, through stacking and destacking – we arrive at design rules including: (i) the oil layer should be as thin as possible to mitigate destacking, while keeping in mind droplet stability and (ii) the droplet should be seated proximal to the injection region at the head of the gel sieving matrix, again to minimize destacking. For example, reducing the droplet size or the thickness of the oil layer, and/or moving the droplet from Δ*x* = 2.5µm to Δ*x* = –2.5 µm, will induce longer retention time and peak broadening during PAGE (**Figure 3d**, **Figure S4, Figure S5, Figure S6, Figure S7**).

To experimentally validate the design rules and optimize the performance of the electroinjection and subsequent PAGE separation, we experimentally scrutinized 45-µm diameter BSA-containing droplets seated in 50-µm diameter microwells. Tilting the assembly allowed us to position each droplet adjacent to the gel lip on the rightmost side of the microwell, thus minimizing the injection region length. Once the droplets were positioned in the microwells, an electric field was applied (*E* = 40 V/cm) and BSA was observed electromigrating out of the droplet, through the minimal injection region, and then into the PAGE sieving matrix. Across a range of droplet positions relative to microwell center and lip, we observed impacts on electromigration and injection dispersion (**Figure 3e**). The observed behaviors were dependent on the droplet oil-layer thickness in the injection region. Once in the sieving matrix, the migration and dispersion of the BSA peak were proportional to the elapsed PAGE time and applied electric field strength (**Figure 3f-h**). At the completion of PAGE and photocapture of each protein peak to the gel matrix, we observed by brightfield microscopy that the originating droplets – now devoid of fluorescently labeled BSA tracer – remained intact in each microwell, thus suggesting robust operation of the selected droplet formulation and overall handling approach under the mechanical and electrical requirements of DropBlot.

We next sought to assess PAGE protein separation performance after electrotransfer of protein sample from the microwell-encapsulated W/O droplets using a soluble, fluorescently labeled protein ladder (OVA, 43 kDa, *D*_inj_ = 3.23 µm^2^/s; *D*_gel_ = 4.65 µm^2^/s; BSA, 66 kDa, *D*_inj_ = 2.40 µm^2^/s, *D*_gel_ = 3.45 µm^2^/s; **Figure 4a-c**). By both simulation and experiment, we observed full resolution of OVA from BSA in 30 s (8%T, 3.3%C gel; *E* = 40 V/cm). The separation resolution (*R_s_* = [Δx / (0.5 × (4σ_1_ + 4σ_2_))] where Δx is peak-to-peak displacement and 4σ is the width of neighboring peaks 1 and 2, respectively) was proportional to the electric field strength and elapsed separation time, as expected, with a slightly higher *R_s_* predicted by simulation (*R_s_* = 4.9) than observed by experiment (*R_s_*= 3.8). Experiments report electrophoretic mobilities of = 3590 µm^2^/(V·s) and = 5830 µm^2^/(V·s) (**Figure S3**). Electroinjection of a large protein species (fluorescently labeled IgG, 150 kDa) was also feasible, as was resolution of the ladder BSA and IgG (**Figure 4d-f**).

**Figure 4.**
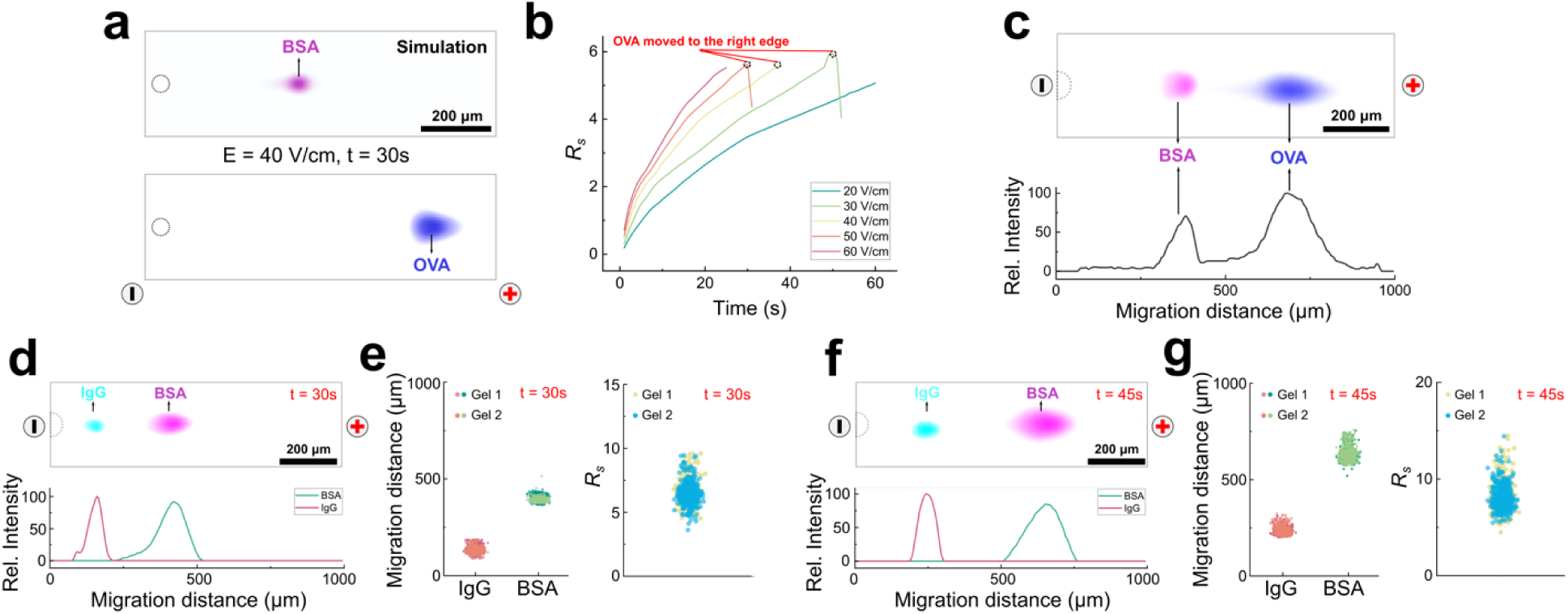
DropBlot assay development utilizing a purified protein ladder. (a) Simulation of PAGE electromigration for BSA and OVA (Δt_PAGE_ = 30 s; *E* = 40 V/cm). (b) Simulation of separation resolution (*R_s_*) of BSA and OVA as a function of applied *E*. Open circles indicate when OVA peak reaches terminus of PAGE separation lane. (c) Fluorescence micrograph of PAGE electromigration for AF555-BSA (pink) and AF647-OVA (blue) (Δt_PAGE_ = 30 s; *E* = 40 V/cm) above fluorescence intensity profile. (d) Fluorescence micrograph of PAGE electromigration for AF555-BSA (pink) and AF488-IgG (blue) (Δt_PAGE_ = 30 s; *E* = 40 V/cm) above fluorescence intensity profile. (e) Comparison of migration distance (left) and separation resolution (right) for PAGE of AF55-BSA and AF488-IgG ladder proteins (n = 500 PAGE lanes; N = 2 devices; Δt_PAGE_ = 30 s; *E* = 40 V/cm). (f) Fluorescence micrograph of PAGE electromigration for AF555-BSA (pink) and AF488-IgG (blue) (Δt_PAGE_ = 45 s; *E* = 40 V/cm) above fluorescence intensity profile. (g) Comparison of migration distance (left) and separation resolution (right) for PAGE of AF55-BSA and AF488-IgG ladder proteins (n = 500 PAGE lanes; N = 2 devices; Δt_PAGE_ = 45 s; *E* = 40 V/cm). For all results: Ø_microwell_ = 50 µm; Ø_droplet_ = 45 µm; Right-aligned droplet (Δx = +2.5 µm) configuration.

### DropBlot analysis of fresh (not fixed) cancer cells

We next sought to scrutinize preparation and analysis of unfixed cells, testing the sample preparation and integration functions of DropBlot. Towards assessing the relationship between antigen-retrieval buffer formulation (including SDS) and the immunoreactivity of any retrieved antigen, we scrutinized fresh cells from well-studied cultured cell lines and the protein targets epithelial cellular adhesion molecule (EpCAM, 35-40 kDa) and intermediate filament protein (VIM, 57 kDa). We make the assumptions that SDS binds proteins with a constant mass ratio (i.e., 1.4 to 1 (SDS: protein)^47^ or 3 to 1 (SDS: protein)^48^), each 45-µm-diameter droplet contains ∼240 fg of SDS, and that each mammalian cell contains ∼100 fg of protein. We reach a working conclusion that the selected droplet volume (Ø = 45 µm, volume = 382 pL) and the selected antigen-retrieval buffer (0.5% (w/v) SDS) provide a mass of SDS sufficient to coat all protein molecules from each single cell. As shown in **Figure 5**, solubilization, electrotransfer, and PAGE analysis were demonstrated using this combination. Immunoreactivity was sufficiently recovered for each protein target, based on successful endpoint immunoblotting.

**Figure 5.**
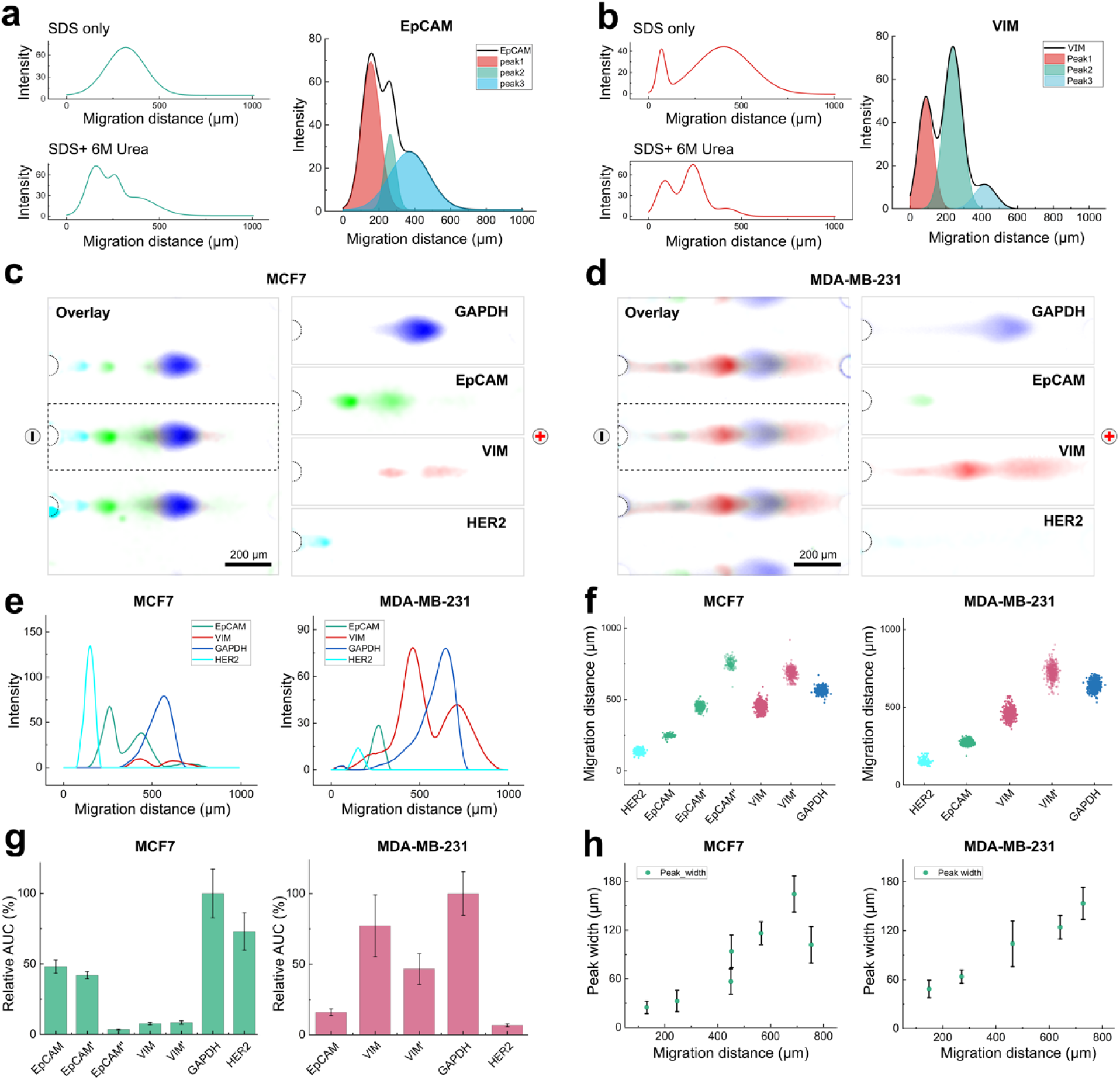
DropBlot assay development utilizing single cells from two unfixed breast cancer cell lines, MCF7 and MDA-MB-231. (a) Fluorescence intensity profile for representative EpCAM PAGE separations from two unfixed MCF7 cells lysed with 0.5% SDS antigen-retrieval buffer without (top) and with (bottom) a 6M urea supplement (Δt_PAGE_ = 30 s; *E* = 40 V/cm). Fluorescence intensity plot (right) shows Gaussian fitting assuming three overlapping EpCAM peaks in SDS+ 6M urea antigen-retrieval buffer formulation. (b) Fluorescence intensity profile for representative VIM PAGE separations from two unfixed MDA-MB-231 cells lysed with 0.5% SDS antigen-retrieval buffer without (top) and with (bottom) a 6M urea supplement (Δt_PAGE_ = 30 s; *E* = 40 V/cm). Fluorescence intensity plot (right) shows Gaussian fitting assuming three overlapping VIM peaks in SDS+ 6M urea antigen-retrieval buffer formulation. Fluorescence micrographs for single-cell PAGE of (c) unfixed MCF7 and (d) unfixed MDA-MB-231 cells for the protein targets EpCAM (green), mesenchymal marker VIM (red), human epidermal growth factor receptor 2 (HER2, cyan), and glycolytic enzyme GAPDH (blue) (Δt_PAGE_ = 30 s; *E* = 60 V/cm). (e) Fluorescence intensity profiles for unfixed cells analyzed in (c) and (d). (f) Migration distance analysis of respective protein targets from single-cell PAGE analysis of MCF7 (left) and MDA-MB-231 cells (right) (n =1000 cells; Δt_PAGE_ = 30 s; *E* = 60 V/cm). (g) Fluorescence area under curve (AUC) for PAGE analyses from (f). (h) PAGE migration distance and peak width for PAGE analyses from (f).

Expanding the repertoire of cell preparation conditions accessible with DropBlot, we explored the possibility of including 6 M urea in the 0.5% (w/v) SDS antigen-retrieval buffer (**Figure 5a-b, Figure S9**). Urea, a strong chaotropic agent, can break hydrogen bonds and unfold hydrophobic protein regions by disrupting hydrophobic interactions^49^. SDS-based antigen-retrieval buffer supplemented with a high concentration of urea (e.g., 6 M) can solubilize a variety of proteoforms by reducing detergent micelles and breaking detergent-protein complexes^50, 51^. We observed that an antigen-retrieval buffer (0.5% (w/v) SDS) supplemented with 6 M urea can resolve additional EpCAM and VIM proteoforms during PAGE of single unfixed cells, with no obvious detrimental effect of the urea supplement on droplet stability (**Figure S10**).

After establishing the DropBlot system as suitable for analysis of protein targets from unfixed single cells, we analyzed our multiplexed cancer-protein panel by immunoblotting the targets EpCAM, VIM, endogenous protein GAPDH, and human epidermal growth factor receptor 2 (HER2). In both a human breast epithelial line (MCF7) and a triple-negative breast cancer line (MDA-MB-231), DropBlot successfully completed unfixed cell and protein sample preparation, then supported immunoblotting of the four targets as shown in **Figure 5c-d**.

Consistent with previous research, we observed that the epithelial cell line, MCF7, had high expression of EpCAM and HER2 and low expression of VIM, relative to a mesenchymal cell line (MDA-MB-231), which had a high expression of VIM and low expression of EpCAM and HER2. The epithelial-to-mesenchymal transition (EMT) can alter the expression of EpCAM, VIM, and HER2^52, 53^. We further observed three resolved EpCAM peaks in the lysate of single MCF7 cells, while MDA-MB-231 cells exhibited one detectable peak. We attribute the cell-line-dependent EpCAM expression to different proteoforms of EpCAM (**Table S2**).

## DropBlot analysis of fixed cancer cells

We scrutinized the DropBlot technology for retrieval and analysis of cancer-related proteins from two types of fixed cancer cells: paraformaldehyde (PFA) and methanol (see **Methods**). We scrutinized PFA and methanol as fixation chemistries, because the two chemistries offer different chemical fixation mechanisms (**Table S3)**. PFA induces covalent bonds between molecules, effectively binding the molecules together to form an insoluble network that changes the mechanical characteristics of the cell surface^54^. In contrast, methanol is understood to denature and precipitate proteins without the formation of covalent bonds^55^.

With these different fixation options in mind, we started by encapsulating fixed cells from each of the well-studied cell lines in the W/O droplets. Each cell was incubated in a droplet with antigen-retrieval buffer at 98°C for 60-120 min. After being subjected to the full DropBlot workflow, immunoblotting of the single fixed-cell lysates showed that protein targets are detectable from PFA-fixed cancer cell lines (MCF7 and MDA-MB-231, fixation time: 15 min). During PAGE, we further observed protein electromigration as being ∼50% slower for targets retrieved from PFA-fixed cells as compared to the respective fresh cancer cell lines (**Figure 6a-b),** as makes sense given what is known about the PFA fixation mechanism. For DropBlot analysis of PFA-fixed cells, we explored the impact of increasing the fixation time (15 vs. 30 min) and increasing the antigen-retrieval incubation time (120 min) and observed that increasing the fixation time (30 min) dramatically reduced the number and intensity of detectable protein peaks **(Figure 6c-e, Figure S11)**. Extended PFA fixation can increase the crosslinking strength between proteins and lipids, making antigen retrieval difficult^56^.

**Figure 6.**
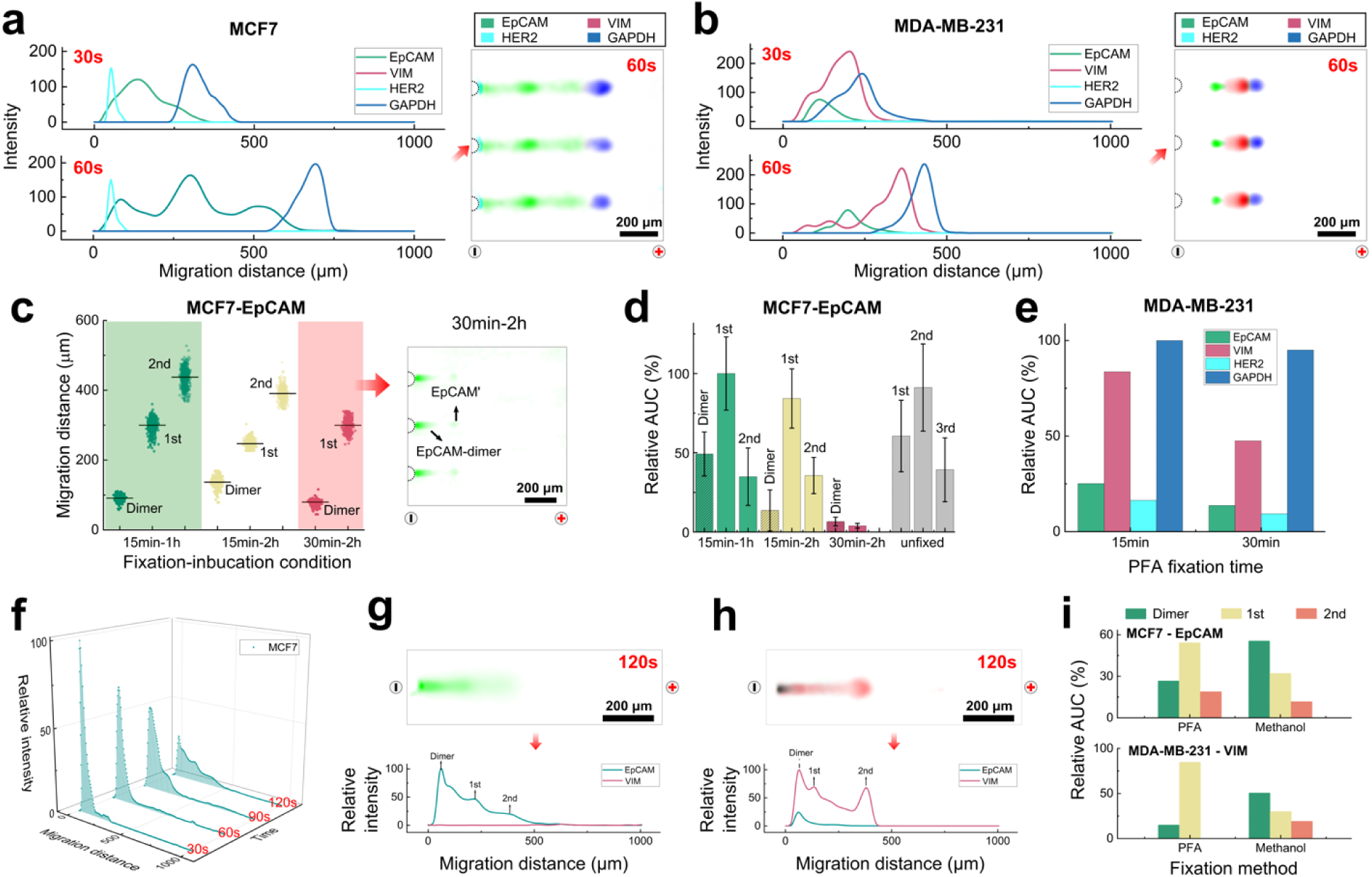
DropBlot assay development utilizing single PFA– and methanol-fixed cancer cells. (a) Fluorescence intensity plots and companion fluorescence micrographs of DropBlot analysis of PFA-fixed MCF7 cells. (b) Same as (a) for PFA-fixed MDA-MB-231 cells. (c) Single-cell western blot EpCAM migration distance for three fixation-incubation conditions (Δt_fixation_ = 15 or 30 min; Δt_incubation_ = 1.0 or 2.0 h at 98°C) with representative fluorescence micrographs from three PFA-fixed MCF7 cells treated with Δt_fixation_ = 30 min; Δt_incubation_ = 2.0 h at 98°C. (d) of single-cell western blot protein peaks for PFA-fixed MCF7 cells (Δt_fixation_ = 15 or 30 min). (e) Same as (d) but for PFA-fixed MDA-MB-231 cells (Δt_fixation_ = 15 or 30 min). (f) Fluorescence intensity from single-cell western blots of EpCAM from methanol-fixed MCF7 cells for Δt_PAGE_ = 30, 60, 90, and 120 s. (g) Fluorescence micrograph from single-cell western blot of EpCAM and VIM from methanol-fixed MCF7 cell above fluorescence intensity profile. (h) Same as (g) for a methanol-fixed MDA-MB-231 cell. (i) Relative AUC of single-cell western blot protein peaks for both cell types and fixation conditions. PFA conditions: Δt_fixation_ = 15 min; Δt_incubation_ = 1.0 h at 98°C; Δt_PAGE_ = 60 s. Methanol conditions: Δt_fixation_ = 15 min; Δt_incubation_ = 1.0 h at 98°C; Δt_PAGE_ = 120 s. For all results, PAGE *E* = 60 V/cm.

Similarly, we scrutinized methanol-fixed MCF7 cells and observed that, while EpCAM was detectable by single-cell immunoblotting, electromigration was even slower than the PFA-fixed cells (**Figure 6f**). We further observed elevated protein signals near the microwells (**Figure 6g-h**, **Figure S12**), suggesting the presence of large protein dimer molecules or even protein aggregates.

We further compared the area under the curve (AUC) of EpCAM and VIM in both MCF7 and MDA-MB-231 cells, using both PFA– and methanol-fixation chemistries. **Figure 6i** illustrates the distribution of EpCAM and VIM proteoforms. In PFA-fixed MCF7 cells, the protein expression profile consisted of: ∼26.7% EpCAM-dimer, 54.3% EpCAM’, and 19.0% EpCAM’’ proteoforms. The protein expression profile of methanol-fixed MCF7 cells consisted of: ∼55.6% EpCAM-dimer, 32.2% EpCAM’, and 11.8% EpCAM’’ proteoforms. PFA-fixed MDA-MB-231 cells contained ∼15.2% VIM-dimer and 84.8% VIM’, whereas methanol-fixed MDA-MB-231 cells showed ∼50.7% VIM-dimer, 30.0% VIM’, and 19.3% VIM’’ proteoforms. The electrotransfer efficiency, defined as the percentage of the protein target successfully extracted and analyzed by DropBlot, spanned a range from 73.2% to 84.8% in PFA-fixed cells, and a range of 44.4% to 49.3% when in methanol-fixed cells. While DropBlot is suitable for analysis of protein targets from both PFA– and methanol-fixed cells, further optimization may be useful to reduce insoluble protein targets retrieved from methanol-fixed cells.

In addition to heat-induced antigen retrieval explored in this first DropBlot report, alternative methods such as enzymatic antigen retrieval (i.e., trypsin, proteinase K, and pepsin) were surveyed in DropBlot. Enzymatic antigen retrieval uses enzymes to digest and break down cross-linked proteins, thus allowing for the recovery of masked protein epitopes that may be inaccessible under standard fixation conditions^57^. We conducted experiments using these enzymatic methods on a PFA-fixed MCF7 cell line and explored various parameters, including enzyme incubation time (5 to 30 min), incubation temperature (room temperature to 37°C), and lysis temperature (room temperature to 100°C) (**Table S4**). Through iterative optimization, we identified pepsin as promising in enzymatic antigen retrieval with separation and detection of the VIM and HER2 proteins from the PFA-fixed MCF7 cells. However, weak or no signals were observed for EpCAM and associated isoforms in any enzymatic antigen-retrieval method explored here (**Figure S13**). This lack of signal could be attributed to two factors: (1) damage to the EpCAM surface protein during enzymatic digestion and/or (2) insufficient cell lysis using an antigen-retrieval buffer composed of 0.5% (w/v) SDS and 6 M urea. Regarding possibility (1): during the enzymatic digestion process, surface proteins, including EpCAM and putative isoforms, may undergo degradation or structural alterations, leading to a loss or reduction in immunoreactivity^58^. The reduction in immunoreactivity may arise from the enzymatic action, which can disrupt the conformation of epitopes or cause proteolytic degradation of the target proteins. As a consequence, the antibody probes used for detection may not recognize the modified or degraded epitopes, resulting in weak or no signals. Regarding possibility (2): the efficiency of cell lysis is crucial for antigen retrieval methods, as cell lysis efficiency affects the release of target proteins and immunoreactivity. The antigen-retrieval buffer used in our experiments (0.5% (w/v) SDS, 6 M urea), may not complete cell lysis and efficient release of the target proteins. Based on these survey results, enzymatic antigen retrieval is a topic of future study for DropBlot.

### DropBlot analysis of fixed clinical specimens

After assay development, we applied DropBlot to protein-target analysis of single, fixed cells dissociated from 11 solid breast tumor specimens. These human-derived tumor tissues were archived for >6 yrs. stored under –80°C conditions without chemical fixation. Prior to scrutiny with DropBlot, the patient-derived cells were thawed and then treated with a range of chemical fixation agents, including PFA and methanol (**Figure 7a**, **Table S5**). After tissue dissociation and PFA-fixation (see Methods), we analyzed protein targets corresponding to epithelial-to-mesenchymal transition (EMT) and tumor cell growth at single-cell level. These targets were chosen because the tumor cells with the mesenchymal phenotype and high expression of HER2 tend to be more aggressive^59, 60^. DropBlot allowed antigen retrieval from 5 fixed-cell specimens, with EpCAM, VIM, and HER2 exhibiting heterogeneous cell-to-cell expression (**Figure 7b-c**). DropBlot analysis showed two VIM proteoforms (VIM’, VIM’’). Previous research has shown the 55-kDa intermediate filament as the most prevalent form of VIM^61^, with truncated VIM generated during cancer metastasis^62–64^.

**Figure 7.**
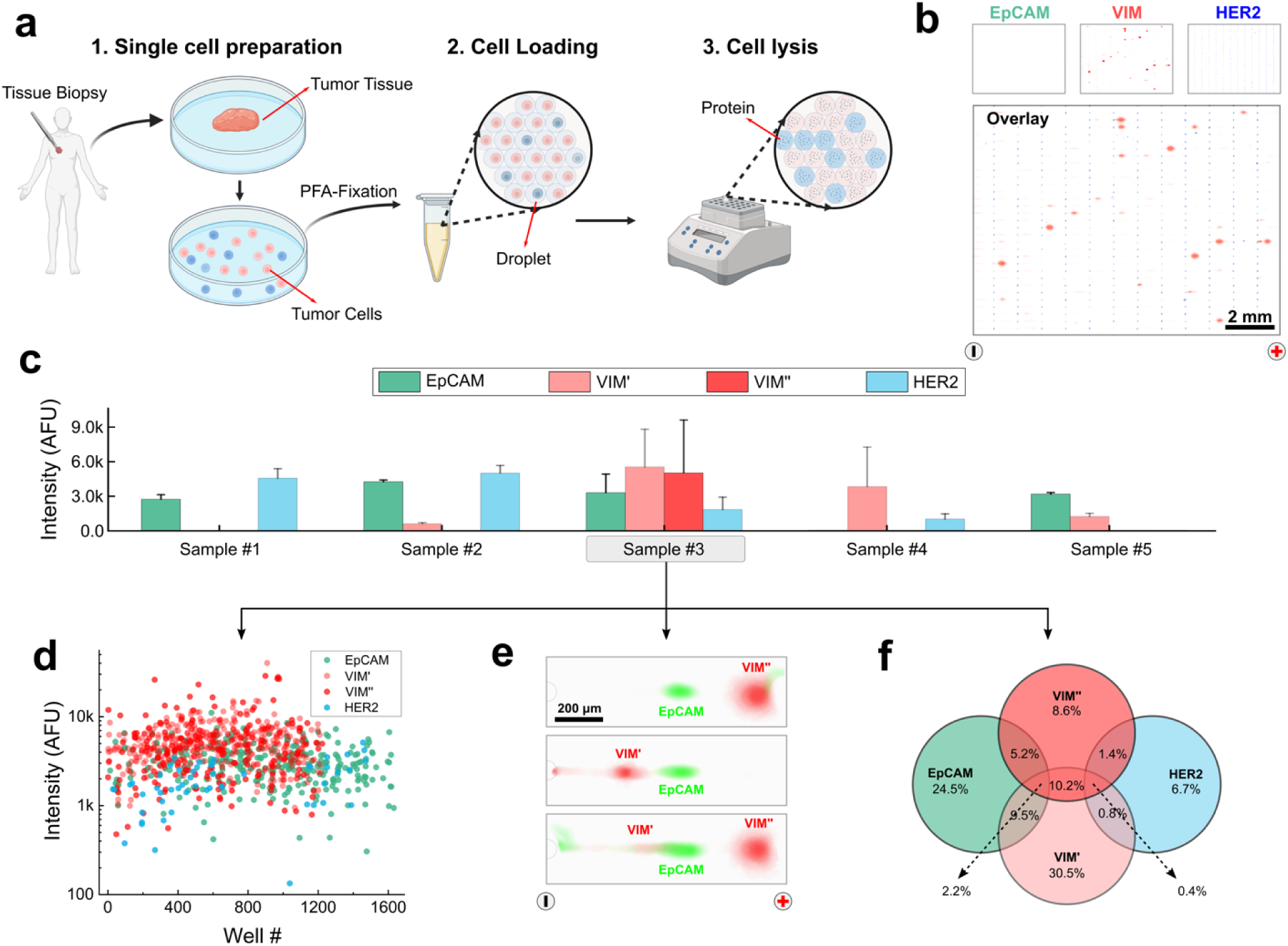
DropBlot analysis of single PFA– and methanol-fixed patient-derived dissociated cancer cells. (a) Schematic of clinical sample preparation workflow for DropBlot. PFA conditions: Δt_fixation_ = 15 min; Δt_incubation_ = 1.0 h at 98°C; Figures generated with BioRender. (b) Fluorescence micrographs of single-cell western blots of EpCAM (Green, AF488-labeled secondary antibody), VIM (Red, AF594-labeled secondary antibody), and HER2 (Blue, AF647-labeled secondary antibody) from PFA-fixed tumor cell. Tumor was classified as triple-positive breast cancer. (c) Mean fluorescence intensity of single-cell western blot analyses of PFA-fixed, patient-derived cells for EpCAM, VIM proteoforms (VIM’, VIM’’), and HER2. Samples #1-2 were fresh cell suspensions. Sample #3-5 were fresh dissociated tissues. (d) Fluorescence AUC of single-cell western blot analyses of PFA-cells in Sample #3 from (c). (e) Fluorescence micrographs of 715 single-cell western blot analyses of PFA-fixed cells from Sample #3 from (c) with EpCAM (Green, AF488-labeled secondary antibody) and VIM (Red, AF594-labeled secondary antibody). (f) Venn diagram reporting the single-cell target-expression profile for each of 715 single PFA-fixed cells from Sample #3 in (c).

In this small cohort, we observed notable patient-to-patient heterogeneity in expression of the cancer protein panel. For example, while Patient #2 displayed high expression of EpCAM and HER2, low expression of VIM’, and no expression of VIM’’, Patient #4 had high expression of VIM’, low expression of HER2, and no detectable expression of EpCAM and VIM’’. Patient #3 with an invasive ductal breast tumor exhibited two VIM proteoforms, which are of interest in relation to increased cancer invasiveness. Two specimens classified as HER2-negative (Patients #1 and #3) exhibited detectable levels of the HER2 protein because –-while not upregulated –-HER2 is nevertheless expected to be present at a baseline level^60^.

In addition to reporting patient-average expression levels of protein targets, DropBlot was employed to provide single-cell level protein expression. We found that the combination and expression level of EpCAM, VIM’, VIM’’, and HER2 were also heterogeneous in the clinical samples (**Figure 7d-e**). **Figure 7f** shows the Venn diagram depicting the distribution of protein-target expression with EpCAM+ alone: 24.5%; VIM’+ alone: 30.5%, VIM’’+ alone: 8.6%; HER2+ alone: 6.7%; EpCAM+/VIM’’: 5.2%; EpCAM+/VIM’: 9.5%; HER2+/VIM’’: 1.4%; HER2+/VIM’: 0.8%; VIM’/VIM’’: 10.2%EpCAM+/VIM’/VIM’’: 2.2%; HER2+/VIM’/VIM’’: 0.4%. Such direct measurement of cancer biomarker distributions –– at the endogenous protein level –-could be important to developing guidance for cancer diagnosis and prognosis.

Based on these results and demonstrated sufficient performance for PFA– and methanol-fixed cells, we see the potential to further mature and refine DropBlot, including exploring additional sample preparation conditions to determine if additional fixation chemistries are compatible with the sample preparation-to-analysis workflow. We are specifically interested in developing DropBlot protocols for the analysis of target antigens in formalin-fixed, paraffin-embedded (FFPE) cell specimens that were not included at a substantial level in this development study. In terms of system design, while DropBlot is utilized here for serial, complementary unit functions (sample preparation to analysis or analysis to detection), we see the modular design as potentially quite powerful for multi-omics questions where coupling of different, complementary analysis stages performed on the same individual cell may lead to breakthroughs in understanding.

## Materials and Methods

### Modeling and simulation

Electric field and protein electromigration were simulated using finite-element modeling of electric current and diluted species transport with COMSOL Multiphysics (version 5.5, COMSOL Inc., Sweden). The simulation geometry for the DropBlot PAGE are presented in the top view shown in **Figure 4a**. Diffusion coefficients in free solution and within the oil layer were calculated using the Stokes-Einstein equation^65^, while the diffusion coefficient in the gel layer was estimated based on published methods^66^. A temperature of 23°C was used as a baseline in all simulations. The initial concentration of protein in the droplet region was 2 µM, while the initial concentration elsewhere in the system was 0 µM. The model was meshed with a Free Triangle Mesh, and we employed a user-controlled override with a maximum element size of 2 µm and a minimum element size of 0.01 µm in the oil layer regions to provide sufficient mesh density in the narrow region.

### Design and fabrication of droplet-generation chip

Single-emulsion droplet generation chips were designed with AutoCAD 2021 (Autodesk, San Francisco, CA). SU-8 3050 (Kayaku Advanced Materials, Westborough, MA) was used to fabricate masters with a height of 60µm following the manufacturer’s instructions. PDMS was prepared using a Sylgard 184 Silicone Elastomer Kit (Ellsworth Adhesives, Germantown, WI) and mixed at a ratio of 10:1. After curing (70°C, 3 hr), the PDMS device was baked for 48 hr at 80°C to recover hydrophobic characteristics.

### Design and fabrication of open-fluidic single-cell western blotting chip

Wafer microfabrication and silanization follow our previous work^67^. Each microwell in the array of 5,000 microwells had a diameter of 50 µm and a depth of 60 µm. Microwell-to-microwell spacing was 1000 µm in the X direction and 300 µm in the Y direction. Fabrication of the polyacrylamide gel layer is based on a developed protocol^68^. To prepare an 8%T polyacrylamide gel, the gel precursor solution was mixed with 30% (w/w) acrylamide/bis-acrylamide (Sigma-Aldrich, St. Louis, MO), N-(3-((3-benzoylphenyl) formamido)propyl) methacrylamide (BPMA, PharmAgra Labs, Brevard, NC), 10× Tris-glycine buffer (Sigma-Aldrich, St. Louis, MO) and ddH_2_O (Sigma-Aldrich, St. Louis, MO). Gels were chemically polymerized for 20 min with 0.08% (w/v) ammonium persulfate (APS, Sigma-Aldrich, St. Louis, MO) and 0.08% (v/v) TEMED (Sigma-Aldrich, St. Louis, MO). After polymerization, gels were carefully released from the wafer by delaminating with a razor blade, and stored in DI water.

### Fluidic generation of W/O droplets for the cell-preparation step

Fresh cells, fixed cells, and purified proteins were resuspended in PBS and injected into the cell inlet of the droplet-generation device (**Figure S1**). Cell suspension and antigen-retrieval buffer were mixed prior to the emulsion region. 1-2% (v/v) Span® 80 surfactant^69^ (Sigma Aldrich, St. Louis, MO) was spiked into mineral oil (Sigma Aldrich, St. Louis, MO) and used as the carrier solution. Flow rates of the substrate and carrier solution were actively controlled by a syringe pump (Chemyx, Stafford, TX). The flow rates of each solution were adjusted to generate droplets of the desired diameter. Droplets were collected in a 1.5 mL Eppendorf® tube or directly loaded onto the top surface of the open-fluidic single-cell western blotting PA gel using gravity with a customized PDMS droplet-delivery channel and holder (**Figure S2**)

### Droplet-stability assays for the cell preparation step

By visual inspection, a suspension of stable droplets will maintain two visible layers (top layer: mineral oil, bottom layer: droplets). Once droplet breakage occurs, three layers develop and equilibrate in the collection tube (top layer: mineral oil, middle layer: droplets, bottom layer: antigen-retrieval buffer). Fluorescently labeled protein targets were diluted to 5 µM with PBS, and included Alexa-Fluor 488 labeled immunoglobulin (IgG, Thermo Fisher Scientific, Waltham, MA), Alexa-Fluor 555 labeled Bovine Serum Albumin (BSA, Thermo Fisher Scientific, Waltham, MA), Alexa-Fluor labeled Ovalbumin 647 (OVA, Thermo Fisher Scientific, Waltham, MA), and rTurboGFP (GFP, Evrogen, Russia). Proteins were encapsulated into 50 µm droplets and observed under an inverted microscope (Olympus IX51, Tokyo, Japan) with a CoolSNAP_HQ2_ camera (Photometrics, Tucson, AZ) for 180 min. Fluorescence images were collected every 3 min, with timing controlled by a mechanical shutter.

### Integration of droplets for the cell preparation step with the single-cell western blot for the cell analysis step

BSA, OVA, and/or IgG proteins were diluted to 0.1 mg/mL and encapsulated in 45-µm droplets containing 0.5% (w/v) SDS. Protein-laden droplets were gravity-settled onto the PA-gel (8%T) surface, which was placed in a customized PAGE chamber (Figure S2, width: 5 cm). A 12.5 mL aliquot of running buffer (1× Tris-glycine and 0.5% (v/v), sodium dodecyl sulfate (SDS, Sigma Aldrich, St. Louis, MO)), was poured onto the chamber. For PAGE, a constant voltage of was applied (200-300 V to obtain E = 40-60 V/cm) using a DC power supply (Bio-Rad PowerPac Basic, Hercules, CA). At PAGE completion, the applied voltage was set to zero and the protein peaks were photo-captured into the PA gel by applying a 45-s UV exposure (Hamamatsu Lightingcure LC5 UV source, HAMAMATSU PHOTONICS, Japan). The chips were rinsed with DI water and imaged with an inverted fluorescence microscope and Genepix® microarray scanner (4300A, Molecular Devices, San Jose, CA). Exposure time and laser power were held constant for all experiments. Determination of the lower limit of detection (LOD) was calculated based on the amount of protein encapsulated by the droplet and droplet volume.

### Culture of cancer cell lines

Human breast cancer cell lines, including MCF7, MCF7/GFP, and MDA-MB-231/GFP (ATCC, Manassas, VA) were cultured in DMEM medium (Thermo Fisher Scientific, Waltham, MA) supplemented with 10% (v/v) fetal bovine serum (GeminiBio, West Sacramento, CA), 1% (v/v) penicillin/streptomycin solution (Thermo Fisher Scientific, Waltham, MA), and 0.1 mM non-essential amino acid solution (Thermo Fisher Scientific, Waltham, MA) in an incubator (37°C, 5% CO2). Prior to DropBlot analysis, adherent cells were released through incubation with 0.05% trypsin-EDTA solution (Thermo Fisher Scientific, Waltham, MA). The concentration of harvested cells was measured with a hemocytometer (Hausser Scientific, Horsham, PA) and resuspended with PBS (Thermo Fisher Scientific, Waltham, MA) to the desired concentration.

### Application of DropBlot to fresh and fixed cells

The cancer cell lines MCF7 and MDA-MB-231 were resuspended with PBS to a concentration of 5 × 10^6^ cells/mL. The antigen-retrieval buffer used for live-cell lysis was 2× Tris-glycine buffer (Sigma Aldrich, St. Louis, MO) supplemented with 1% SDS and 12 M urea (Sigma Aldrich, St. Louis, MO). Flow rates of cell solution, antigen-retrieval buffer, and carrier layer (oil) were 1.0, 1.0, 12.0 µL/min to generate droplets with diameters of ∼45 µm. The final concentration of antigen-retrieval buffer was 1× Tris-glycine supplemented with 0.5% SDS and 6 M urea. Cell-laden droplets were loaded onto the top surface of the single-cell western blotting device (8%T polyacrylamide gel, stippled with microwells). The PDMS slab was then relocated to cover the droplet-filled microwells and excess mineral oil was flushed out with running buffer (1X Tris-glycine supplemented with 1% SDS). Cells were lysed for at least 10 min once encapsulated in droplets at room temperature.

12.5 mL of running buffer was poured into the chamber and protein target electro-transfer and PAGE were initiated by applying a 300V constant voltage for 30 s across the device to achieve an average E = 40 – 60 V/cm across the gel. At PAGE completion, the applied potential was zeroed out, the proteins halted electromigration, and stationary protein peaks were photo-captured to the benzophenone in the PA gel by application of a 45 s pulse of UV light. The gel was then rinsed briefly with deionized water and stored in Tris-buffered saline with Tween 20 (TBST, Cell Signaling Technology, Danvers, MA) overnight to remove excess oil and residual antigen-retrieval buffer.

To generate PFA-fixed cells, the cancer cell lines MCF7 and MDA-MB-231 were fixed with 4% paraformaldehyde (PFA, Alfa Aesar, Haverhill, MA) for 15-30 min at room temperature following the manufacturer’s protocol. After fixation, cells were washed 3× with PBS to remove excess PFA and resuspended with PBS to achieve a concentration of ∼5 × 10^6^ cells/mL. The antigen-retrieval buffer used for live-cell PFA fixation was 2× Tris-glycine buffer supplemented with 2% SDS and 12 M urea. Flow rates of cell solution, antigen-retrieval buffer, and carrier layer (oil) were 0.5, 0.5, and 5 µL/min, respectively, to generate 45-µm diameter droplets. The final concentration of antigen-retrieval buffer was 1× Tris-glycine buffer supplemented with 1% SDS and 6 M urea. The droplets were collected and incubated in a 1.5 mL Eppendorf tube at 98°C for 1 – 2 hr, and then loaded onto the 8%T polyacrylamide gel. Running buffer (12.5 mL) was poured into the chamber and the separation was initiated by supplying 300V constant voltage for an average electric field of 40-60 V/cm across the gel for 30-60 s. After PAGE, the proteins were photo-captured by 45-s UV exposure. The gel was then rinsed briefly with deionized water and stored in TBST.

### For methanol-fixed cells

the cancer cell lines MCF7 and MDA-MB-231 were again used as model cells, now fixed with ice-cold methanol (Sigma Aldrich, St. Louis, MO) for 15 min at – 20°C following the manufacturer’s protocol. After fixation, cells were washed 3× with PBS to remove excess methanol and resuspended with PBS to achieve a concentration of ∼5e6 cells/mL. The live-cell antigen-retrieval buffer was 2× Tris-glycine buffer supplemented with 2% SDS and 12 M urea. Flow rates of the cell solution, antigen-retrieval buffer, and carrier layer (oil) were 0.5, 0.5, and 5 µL/min, respectively, to generate 45-µm diameter droplets. The final concentration of antigen-retrieval buffer was 1× Tris-glycine buffer supplemented with 1% SDS and 6 M urea. The droplets were collected and incubated in a 1.5 mL Eppendorf® tube at 98°C for 1 hour, and then loaded onto 8%T polyacrylamide gel. 12.5 mL of running buffer was poured into the chamber and the separation was initiated by supplying 300 V constant voltage for 30-120 s to reach an average electric field of 40-60 V/cm across the gel. After PAGE, the proteins were photo-captured by applying UV light to the gel for 45 s. The gel was then rinsed briefly with deionized water and stored in TBST.

### Antigen retrieval using enzymatic methods

Cancer cell line, MCF7, was fixed with PFA for 15 min at room temperature. The fixed cells were washed with PBS for three times and aliquoted to a 1.5 mL Eppendorf tube with 1e6 cell/tube. Trypsin antigen retrieval solution (ab970, Abcam, Cambridge, United Kingdom), proteinase K antigen retrieval solution (ab64220), and pepsin antigen retrieval solution (ab64201) were used following the manufacturer’s instructions. The detailed incubation and lysis conditions are listed in **Table S4**. After incubation, cells were encapsulated in 45 µm droplet and the protein lysate was separated by supplying 300V constant voltage across the gel for 30s. After PAGE, the proteins were photo-captured by 45s’ UV exposure. The gel was then rinsed briefly with deionized water and stored in TBST.

### Patient-derived tissue samples from a biospecimen repository

Primary-tumor tissue was obtained anonymized and blinded from Stanford Cancer Institute’s Tissue Procurement Shared Resource facility. Human-derived tumor specimens were archived for > 6 yr prior to DropBlot analysis, stored at –80 °C.

### Fresh tumor specimens (cell suspension or tissue)

Frozen samples were thawed in a water bath at 37°C for 1 min and mixed with 10 mL of DMEM medium. Samples were centrifuged at 300 × g for 5 min to remove the supernatant. For the fresh cell suspension, the samples were fixed with 4% PFA for 15 min and resuspended with PBS to a concentration of 5 × 10^6^ cells/mL. For the fresh tissue (Figure 7a), a 1-g tissue specimen was weighed and placed in a petri dish containing 5 mL of 37°C DMEM medium. Using a scalpel and tweezer, the tissue was coarsely dissected into fragments <0.75 mm in diameter. A tissue suspension was constituted by adding 5 mL of Tumor & Tissue Dissociation Regent (TTDR, BD Bioscience, San Jose, CA), and then incubating the mixture at 37 °C for 30 min with frequent agitation. After incubation, 25 mL of Dulbecco’s Phosphate Buffered Saline (DPBS, Thermo Fisher Scientific, Waltham, MA) containing 1% BSA and 2 mM EDTA (Thermo Fisher Scientific, Waltham, MA) was added.

Large tissue/cell clusters were removed with a 70-µm cell strainer and then centrifuged at 300 x g for 5 min to remove the supernatant. A cell pellet formed and was resuspended in 2 mL of 1× lysis buffer (Tonbo Biosciences, San Diego, CA) and incubated at room temperature for 15 min. A 40 mL aliquot of DPBS containing 1% BSA and 2 mM EDTA was then added to the mixture. After removing the supernatant, the cells were fixed following the fixation protocols described elsewhere. Cells were counted using a hemocytometer and resuspended to 5 × 10^6^ cells/mL with PBS.

### Formalin-fixed, paraffin-embedded (FFPE) tumor specimens

Frozen tissue samples were thawed in water bath at 60°C for 2 hr, and bathed in 10 mL xylene (Sigma Aldrich, St. Louis, MO) for 5 min (twice). The samples were rehydrated with 96% ethanol, 90% ethanol, 70% ethanol, 50% ethanol, and PBS for 5 min, then washed twice. The cells were fixed following the fixation protocols described elsewhere and resuspended to 5 × 10^6^ cells/mL with PBS.

### Immunoprobing and fluorescence imaging for the cell analysis step

The primary antibody immunoprobing solution was prepared by diluting stock solutions of primary antibodies in 2% (w/v) BSA/TBST solution to achieve an antibody concentration of 0.05 µg/µL (single antibody). Primary antibodies used were EpCAM, VIM, HER2, and GAPDH (Abcam, Cambridge, United Kingdom). The single-cell western blotting device (gel slide) was treated with 80 µL of primary antibody immunoprobing solution and incubated at room temperature for 2 hr. After incubation, each single-cell western blotting device was washed twice with TBST buffer for 1 hour. The secondary antibody immunoprobing solution was prepared by diluting stock solutions of primary antibodies in 2% (w/v) BSA/TBST solution to achieve a concentration of 0.05 µg/µL (single antibody). The single-cell western blotting device was incubated with 80 µL of secondary antibody immunoprobing solution at room temperature for 2 hr. After incubation, the single-cell western blotting device was washed twice with TBST buffer for 1 hr. Before fluorescence imaging, the single-cell western blotting device was washed 3× with DI water to remove excess salts, and dried with nitrogen gun. The single-cell western blotting device was imaged with a Genepix Microarray Scanner. Images were analyzed using custom analysis scripts in MATLAB (MathWorks, Natick, MA).

## Acknowledgments

This work was funded by the National Institutes of Health (NIH) R01CA20301. We sincerely thank Herr Lab members Gabriela Lomeli and Ana Martinez for initial training on the operation of the single-cell western blot. We acknowledge all members of the Herr Lab at UC Berkeley. We are grateful to the R&D Machine Shop at UC Berkeley for the fabrication of the chamber and to patients who selflessly donated biospecimens to the Stanford Cancer Institute’s Tissue Procurement Shared Resource facility.

## Author contributions

Y.L. and A.E.H. designed the hybrid microfluidic platform (DropBlot). Y.L. and A.E.H. designed experiments. Y.L. performed DropBlot assay on purified proteins, fresh & fixed cell lines, and fresh & fixed clinical samples. Y.L. designed software and performed data analysis. Y.L. and A.E.H. wrote the manuscript.

## Competing interests

No competing interests.

## Additional information

Supplementary information is available.

## Notes

### Competing Interest Statement

The authors have declared no competing interest.

## References

(1) Blow, N. Tissue issues. Nature 2007, 448 (7156), 959–960.

(2) Hudis, C. A. Trastuzumab—mechanism of action and use in clinical practice. New Engl J Med 2007, 357 (1), 39–51.

(3) Werner, M.; Chott, A.; Fabiano, A.; Battifora, H. Effect of formalin tissue fixation and processing on immunohistochemistry. The American journal of surgical pathology 2000, 24 (7), 1016–1019.

(4) Fujiwara, K. Techniques for localizing contractile proteins with fluorescent antibodies. Curr Top Dev Biol 1980, 14 (Pt 2), 271–296. DOI: 10.1016/s0070-2153(08)60198-2.

(5) Kuzmin, A. N.; Pliss, A.; Prasad, P. N. Changes in biomolecular profile in a single nucleolus during cell fixation. Anal Chem 2014, 86 (21), 10909–10916. DOI: 10.1021/ac503172b.

(6) Hewitt, S. M.; Lewis, F. A.; Cao, Y.; Conrad, R. C.; Cronin, M.; Danenberg, K. D.; Goralski, T. J.; Langmore, J. P.; Raja, R. G.; Williams, P. M. Tissue handling and specimen preparation in surgical pathology: issues concerning the recovery of nucleic acids from formalin-fixed, paraffin-embedded tissue. Archives of pathology & laboratory medicine 2008, 132 (12), 1929–1935.

(7) Sompuram, S. R.; Vani, K.; Messana, E.; Bogen, S. A. A molecular mechanism of formalin fixation and antigen retrieval. American journal of clinical pathology 2004, 121 (2), 190–199.

(8) Ly, A.; Buck, A.; Balluff, B.; Sun, N.; Gorzolka, K.; Feuchtinger, A.; Janssen, K.-P.; Kuppen, P. J.; van de Velde, C. J.; Weirich, G. High-mass-resolution MALDI mass spectrometry imaging of metabolites from formalin-fixed paraffin-embedded tissue. Nature protocols 2016, 11 (8), 1428–1443.

(9) O’Leary, T.; Fowler, C.; Evers, D.; Mason, J. Protein fixation and antigen retrieval: chemical studies. Biotechnic & Histochemistry 2009, 84 (5), 217–221.

(10) Cottu, P. H.; Asselah, J.; Lae, M.; Pierga, J. Y.; Diéras, V.; Mignot, L.; Sigal-Zafrani, B.; Vincent-Salomon, A. Intratumoral heterogeneity of HER2/neu expression and its consequences for the management of advanced breast cancer. Ann Oncol 2008, 19 (3), 595–597. DOI: 10.1093/annonc/mdn021 From NLM.

(11) Mantsiou, A.; Makridakis, M.; Fasoulakis, K.; Katafigiotis, I.; Constantinides, C. A.; Zoidakis, J.; Roubelakis, M. G.; Vlahou, A.; Lygirou, V. Proteomics Analysis of Formalin Fixed Paraffin Embedded Tissues in the Investigation of Prostate Cancer. J Proteome Res 2020, 19 (7), 2631–2642. DOI: 10.1021/acs.jproteome.9b00587.

(12) Ramos-Vara, J. A. Technical aspects of immunohistochemistry. Veterinary pathology 2005, 42 (4), 405–426.

(13) Polak, J. M.; Van Noorden, S. Introduction to immunocytochemistry; BIOS Scientific Publishers Oxford, 1997.

(14) Gown, A. M. Current issues in ER and HER2 testing by IHC in breast cancer. Modern Pathol 2008, 21, S8–S15. DOI: 10.1038/modpathol.2008.34.

(15) Sabbatino, F.; Villani, V.; Yearley, J. H.; Deshpande, V.; Cai, L.; Konstantinidis, I. T.; Moon, C.; Nota, S.; Wang, Y. Y.; Al-Sukaini, A.;, et al. PD-L1 and HLA Class I Antigen Expression and Clinical Course of the Disease in Intrahepatic Cholangiocarcinoma. Clin Cancer Res 2016, 22 (2), 470–478. DOI: 10.1158/1078-0432.Ccr-15-0715.

(16) Irish, J. M.; Myklebust, J. H.; Alizadeh, A. A.; Houot, R.; Sharman, J. P.; Czerwinski, D. K.; Nolan, G. P.; Levy, R. B-cell signaling networks reveal a negative prognostic human lymphoma cell subset that emerges during tumor progression. Proceedings of the National Academy of Sciences 2010, 107 (29), 12747–12754.

(17) Adan, A.; Alizada, G.; Kiraz, Y.; Baran, Y.; Nalbant, A. Flow cytometry: basic principles and applications. Critical reviews in biotechnology 2017, 37 (2), 163–176.

(18) van Remoortere, A.; van Zeijl, R. J. M.; van den Oever, N.; Franck, J.; Longuespee, R.; Wisztorski, M.; Salzet, M.; Deelder, A. M.; Fournier, I.; McDonnell, L. A. MALDI Imaging and Profiling MS of Higher Mass Proteins from Tissue. J Am Soc Mass Spectr 2010, 21 (11), 1922–1929. DOI: 10.1016/j.jasms.2010.07.011.

(19) Zhu, Y.; Clair, G.; Chrisler, W. B.; Shen, Y.; Zhao, R.; Shukla, A. K.; Moore, R. J.; Misra, R. S.; Pryhuber, G. S.; Smith, R. D.;, et al. Proteomic Analysis of Single Mammalian Cells Enabled by Microfluidic Nanodroplet Sample Preparation and Ultrasensitive NanoLC-MS. Angew Chem Int Ed Engl 2018, 57 (38), 12370–12374. DOI: 10.1002/anie.201802843 From NLM Medline.

(20) Jackson, H. W.; Fischer, J. R.; Zanotelli, V. R.; Ali, H. R.; Mechera, R.; Soysal, S. D.; Moch, H.; Muenst, S.; Varga, Z.; Weber, W. P. The single-cell pathology landscape of breast cancer. Nature 2020, 578 (7796), 615–620.

(21) Su, P.; McGee, J. P.; Durbin, K. R.; Hollas, M. A. R.; Yang, M.; Neumann, E. K.; Allen, J. L.; Drown, B. S.; Butun, F. A.; Greer, J. B.;, et al. Highly multiplexed, label-free proteoform imaging of tissues by individual ion mass spectrometry. Sci Adv 2022, 8 (32), eabp9929. DOI: 10.1126/sciadv.abp9929.

(22) Eyer, K.; Doineau, R. C. L.; Castrillon, C. E.; Briseno-Roa, L.; Menrath, V.; Mottet, G.; England, P.; Godina, A.; Brient-Litzler, E.; Nizak, C.;, et al. Single-cell deep phenotyping of IgG-secreting cells for high-resolution immune monitoring. Nat Biotechnol 2017, 35 (10), 977–982. DOI: 10.1038/nbt.3964.

(23) Gebreyesus, S. T.; Siyal, A. A.; Kitata, R. B.; Chen, E. S.; Enkhbayar, B.; Angata, T.; Lin, K. I.; Chen, Y. J.; Tu, H. L. Streamlined single-cell proteomics by an integrated microfluidic chip and data-independent acquisition mass spectrometry. Nat Commun 2022, 13 (1), 37. DOI: 10.1038/s41467-021-27778-4.

(24) Blazek, M.; Santisteban, T. S.; Zengerle, R.; Meier, M. Analysis of fast protein phosphorylation kinetics in single cells on a microfluidic chip. Lab on a Chip 2015, 15 (3), 726–734.

(25) Lu, Y.; Xue, Q.; Eisele, M. R.; Sulistijo, E. S.; Brower, K.; Han, L.; Amir el, A. D.; Pe’er, D.; Miller-Jensen, K.; Fan, R. Highly multiplexed profiling of single-cell effector functions reveals deep functional heterogeneity in response to pathogenic ligands. Proc Natl Acad Sci U S A 2015, 112 (7), E607–615. DOI: 10.1073/pnas.1416756112.

(26) Shembekar, N.; Hu, H.; Eustace, D.; Merten, C. A. Single-cell droplet microfluidic screening for antibodies specifically binding to target cells. Cell reports 2018, 22 (8), 2206–2215.

(27) Liu, Y.; Vieira, R. M. S.; Mao, L. Simultaneous and Multimodal Antigen-Binding Profiling and Isolation of Rare Cells via Quantitative Ferrohydrodynamic Cell Separation. ACS Nano 2023, 17 (1), 94–110. DOI: 10.1021/acsnano.2c04542.

(28) Kurien, B. T.; Scofield, R. H. Western blotting. Methods 2006, 38 (4), 283–293.

(29) Chu, W. S.; Liang, Q.; Liu, J.; Wei, M. Q.; Winters, M.; Liotta, L.; Sandberg, G.; Gong, M. A nondestructive molecule extraction method allowing morphological and molecular analyses using a single tissue section. Lab Invest 2005, 85 (11), 1416–1428. DOI: 10.1038/labinvest.3700337 From NLM Medline.

(30) Becker, K. F.; Schott, C.; Hipp, S.; Metzger, V.; Porschewski, P.; Beck, R.; Nährig, J.; Becker, I.; Höfler, H. Quantitative protein analysis from formalin-fixed tissues: implications for translational clinical research and nanoscale molecular diagnosis. The Journal of Pathology: A Journal of the Pathological Society of Great Britain and Ireland 2007, 211 (3), 370–378.

(31) Grist, S. M.; Mourdoukoutas, A. P.; Herr, A. E. 3D projection electrophoresis for single-cell immunoblotting. Nature communications 2020, 11 (1), 6237.

(32) Hughes, A. J.; Spelke, D. P.; Xu, Z.; Kang, C.-C.; Schaffer, D. V.; Herr, A. E. Single-cell western blotting. Nature methods 2014, 11 (7), 749–755.

(33) Smith, A.; Galli, M.; Piga, I.; Denti, V.; Stella, M.; Chinello, C.; Fusco, N.; Leni, D.; Manzoni, M.; Roversi, G. Molecular signatures of medullary thyroid carcinoma by matrix-assisted laser desorption/ionisation mass spectrometry imaging. Journal of proteomics 2019, 191, 114–123.

(34) Heijs, B.; Holst, S.; Briaire-de Bruijn, I. H.; Van Pelt, G. W.; de Ru, A. H.; van Veelen, P. A.; Drake, R. R.; Mehta, A. S.; Mesker, W. E.; Tollenaar, R. A. Multimodal mass spectrometry imaging of N-glycans and proteins from the same tissue section. Anal Chem 2016, 88 (15), 7745–7753.

(35) Sompuram, S. R.; Vani, K.; Schaedle, A. K.; Balasubramanian, A.; Bogen, S. A. Selecting an Optimal Positive IHC Control for Verifying Antigen Retrieval. J Histochem Cytochem 2019, 67 (4), 275–289. DOI: 10.1369/0022155418824092.

(36) Buczak, K.; Kirkpatrick, J. M.; Truckenmueller, F.; Santinha, D.; Ferreira, L.; Roessler, S.; Singer, S.; Beck, M.; Ori, A. Spatially resolved analysis of FFPE tissue proteomes by quantitative mass spectrometry. Nat Protoc 2020, 15 (9), 2956–2979. DOI: 10.1038/s41596-020-0356-y.

(37) Baret, J. C. Surfactants in droplet-based microfluidics. Lab Chip 2012, 12 (3), 422–433. DOI: 10.1039/c1lc20582j.

(38) Mazutis, L.; Gilbert, J.; Ung, W. L.; Weitz, D. A.; Griffiths, A. D.; Heyman, J. A. Single-cell analysis and sorting using droplet-based microfluidics. Nat Protoc 2013, 8 (5), 870–891. DOI: 10.1038/nprot.2013.046.

(39) Aronson, M. P.; Petko, M. F. Highly concentrated water-in-oil emulsions: Influence of electrolyte on their properties and stability. Journal of colloid and interface science 1993, 159 (1), 134–149.

(40) Villa, C. H.; Lawson, L. B.; Li, Y. M.; Papadopoulos, K. D. Internal coalescence as a mechanism of instability in water-in-oil-in-water double-emulsion globules. Langmuir 2003, 19 (2), 244–249. DOI: 10.1021/la026324d.

(41) Dittrich, P. S.; Jahnz, M.; Schwille, P. A new embedded process for compartmentalized cell-free protein expression and on-line detection in microfluidic devices. Chembiochem 2005, 6 (5), 811–814. DOI: 10.1002/cbic.200400321.

(42) Benson, B. R.; Stone, H. A.; Prud’homme, R. K. An “off-the-shelf” capillary microfluidic device that enables tuning of the droplet breakup regime at constant flow rates. Lab on a Chip 2013, 13 (23), 4507–4511.

(43) Demetriades, K.; McClements, D. J. Influence of sodium dodecyl sulfate on the physicochemical properties of whey protein-stabilized emulsions. Colloid Surface A 2000, 161 (3), 391–400. DOI: Doi 10.1016/S0927-7757(99)00210-1.

(44) De Aguiar, H. B.; Strader, M. L.; de Beer, A. G.; Roke, S. Surface structure of sodium dodecyl sulfate surfactant and oil at the oil-in-water droplet liquid/liquid interface: a manifestation of a nonequilibrium surface state. The journal of physical chemistry B 2011, 115 (12), 2970–2978.

(45) Rouabeh, J.; M’barki, L.; Hammami, A.; Jallouli, I.; Driss, A. Studies of different types of insulating oils and their mixtures as an alternative to mineral oil for cooling power transformers. Heliyon 2019, 5 (3), e01159.

(46) Khademi, M.; Cheng, S. S. Y.; Barz, D. P. J. Charge and Electrical Double Layer Formation in a Nonpolar Solvent Using a Nonionic Surfactant. Langmuir 2020, 36 (19), 5156–5164. DOI: 10.1021/acs.langmuir.0c00311.

(47) Reynolds, J. A.; Tanford, C. Binding of dodecyl sulfate to proteins at high binding ratios. Possible implications for the state of proteins in biological membranes. Proceedings of the National Academy of Sciences 1970, 66 (3), 1002–1007.

(48) Hames, B. D. Gel electrophoresis of proteins: a practical approach; OUP Oxford, 1998.

(49) Rabilloud, T. Solubilization of proteins for electrophoretic analyses. Electrophoresis 1996, 17 (5), 813–829. DOI: 10.1002/elps.1150170503.

(50) Wisniewski, J. R.; Zougman, A.; Nagaraj, N.; Mann, M. Universal sample preparation method for proteome analysis. Nat Methods 2009, 6 (5), 359–362. DOI: 10.1038/nmeth.1322.

(51) Glatter, T.; Ahrne, E.; Schmidt, A. Correction to “Comparison of Different Sample Preparation Protocols Reveals Lysis Buffer-Specific Extraction Biases in Gram-Negative Bacteria and Human Cells”. J Proteome Res 2016, 15 (2), 679. DOI: 10.1021/acs.jproteome.6b00012.

(52) Kalluri, R.; Weinberg, R. A. The basics of epithelial-mesenchymal transition. J Clin Invest 2009, 119 (6), 1420–1428. DOI: 10.1172/JCI39104.

(53) Carpenter, R. L.; Paw, I.; Dewhirst, M. W.; Lo, H.-W. Akt phosphorylates and activates HSF-1 independent of heat shock, leading to Slug overexpression and epithelial–mesenchymal transition (EMT) of HER2-overexpressing breast cancer cells. Oncogene 2015, 34 (5), 546–557.

(54) Kim, S. O.; Kim, J.; Okajima, T.; Cho, N. J. Mechanical properties of paraformaldehyde-treated individual cells investigated by atomic force microscopy and scanning ion conductance microscopy. Nano Converg 2017, 4 (1), 5. DOI: 10.1186/s40580-017-0099-9.

(55) Wang, X.; Yu, L.; Wu, A. R. Correction to: The effect of methanol fixation on single-cell RNA sequencing data. BMC Genomics 2021, 22 (1), 554. DOI: 10.1186/s12864-021-07806-9.

(56) Erturk, A.; Becker, K.; Jahrling, N.; Mauch, C. P.; Hojer, C. D.; Egen, J. G.; Hellal, F.; Bradke, F.; Sheng, M.; Dodt, H. U. Three-dimensional imaging of solvent-cleared organs using 3DISCO. Nat Protoc 2012, 7 (11), 1983–1995. DOI: 10.1038/nprot.2012.119.

(57) Yamashita, S. Heat-induced antigen retrieval: mechanisms and application to histochemistry. Prog Histochem Cytochem 2007, 41 (3), 141–200. DOI: 10.1016/j.proghi.2006.09.001.

(58) Schon, M. P.; Schon, M.; Mattes, M. J.; Stein, R.; Weber, L.; Alberti, S.; Klein, C. E. Biochemical and immunological characterization of the human carcinoma-associated antigen MH 99/KS 1/4. Int J Cancer 1993, 55 (6), 988–995. DOI: 10.1002/ijc.2910550619.

(59) Lambert, A. W.; Pattabiraman, D. R.; Weinberg, R. A. Emerging Biological Principles of Metastasis. Cell 2017, 168 (4), 670–691. DOI: 10.1016/j.cell.2016.11.037.

(60) Loibl, S.; Gianni, L. HER2-positive breast cancer. The Lancet 2017, 389 (10087), 2415–2429.

(61) Rose, M. L. Role of anti-vimentin antibodies in allograft rejection. Hum Immunol 2013, 74 (11), 1459–1462. DOI: 10.1016/j.humimm.2013.06.006.

(62) Whipple, R. A.; Balzer, E. M.; Cho, E. H.; Matrone, M. A.; Yoon, J. R.; Martin, S. S. Vimentin filaments support extension of tubulin-based microtentacles in detached breast tumor cells. Cancer Research 2008, 68 (14), 5678–5688. DOI: 10.1158/0008-5472.Can-07-6589.

(63) Zhang, H.; Wu, X. S.; Xiao, Y. Z.; Wu, L. Q.; Peng, Y.; Tang, W. M.; Liu, G. N.; Sun, Y.; Wang, J.; Zhu, H. Q.;, et al. Coexpression of FOXK1 and vimentin promotes EMT, migration, and invasion in gastric cancer cells. J Mol Med 2019, 97 (2), 163–176. DOI: 10.1007/s00109-018-1720-z.

(64) Makise, M.; Nakamura, H.; Kuniyasu, A. The role of vimentin in the tumor marker Nup88-dependent multinucleated phenotype. Bmc Cancer 2018, 18. DOI: ARTN 519 10.1186/s12885-018-4454-y.

(65) Miller, C. C. The Stokes Einstein law for diffusion in solution. P R Soc Lond a-Conta 1924, 106 (740), 724–749. DOI: DOI 10.1098/rspa.1924.0100.

(66) Park, I. H.; Johnson, C. S.; Gabriel, D. A. Probe Diffusion in Polyacrylamide Gels as Observed by Means of Holographic Relaxation Methods – Search for a Universal Equation. Macromolecules 1990, 23 (5), 1548–1553. DOI: DOI 10.1021/ma00207a052.

(67) Rosàs-Canyelles, E.; Modzelewski, A. J.; Geldert, A.; He, L.; Herr, A. E. Multimodal detection of protein isoforms and nucleic acids from mouse pre-implantation embryos. Nature protocols 2021, 16 (2), 1062–1088.

(68) Kang, C. C.; Yamauchi, K. A.; Vlassakis, J.; Sinkala, E.; Duncombe, T. A.; Herr, A. E. Single cell-resolution western blotting. Nat Protoc 2016, 11 (8), 1508–1530. DOI: 10.1038/nprot.2016.089 From NLM Medline.

(69) Jiao, J.; Burgess, D. J. Rheology and stability of water-in-oil-in-water multiple emulsions containing Span 83 and Tween 80. Aaps Pharmsci 2003, 5, 62–73.

