## Supplementary information for "DropBlot: single-cell western blotting of chemically fixed cancer cells"

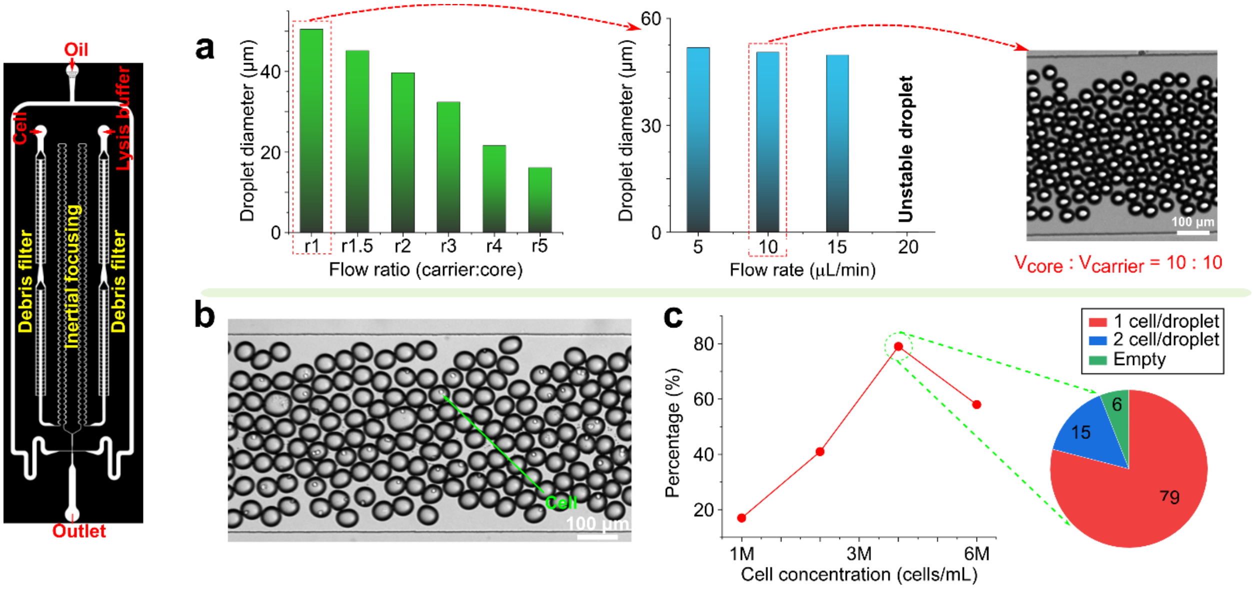


**Figure S1.** Optimization of droplet generation for cell encapsulation. (a) Study of droplet size with different parameters: flow ratio between carrier and core medium (left); flow rate of core medium when the flow ratio was 1 (middle). Stable droplets of 50 µm in diameter were generated (right). (b) Bright-field images of cell encapsulation when the initial concentration was 3 M ($3.0\times10^{6}$) cells/mL. (c) The percentage of droplets containing one cell with a different initial cell concentration.

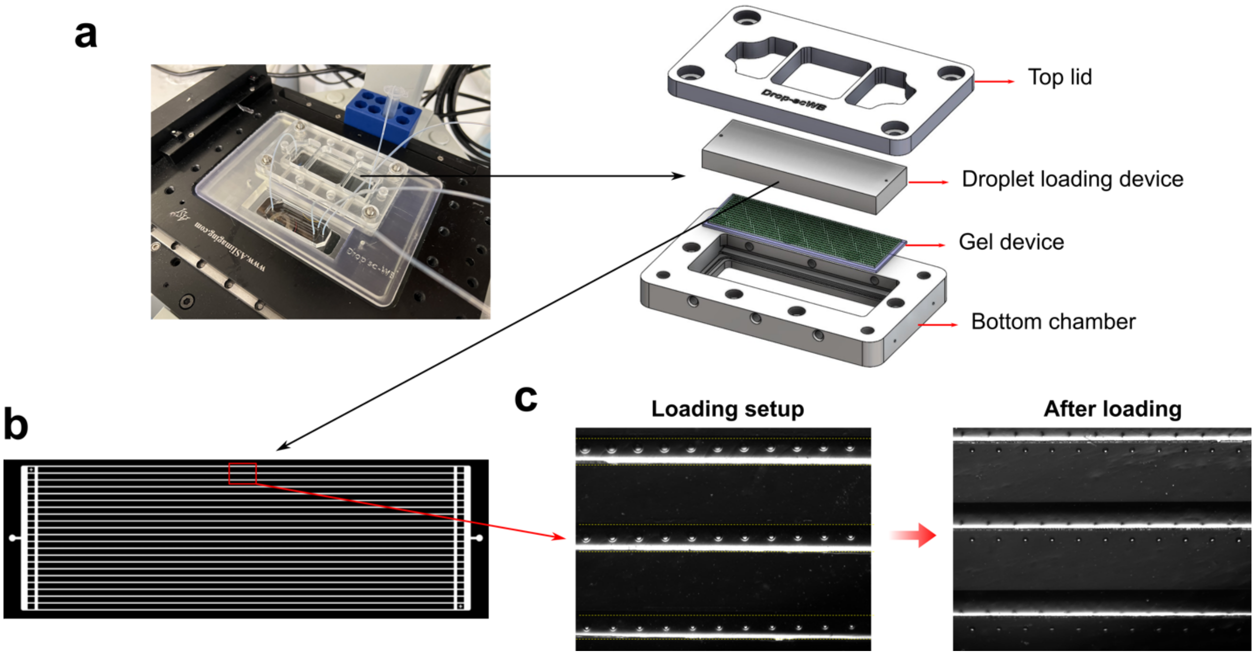


**Figure S2.** All-in-one reaction holder setup and assembly. (a) overview (left) and components (right) of the all-in-one reaction holder. The top lid and bottom chamber are made of PMMA. PA-Gel device and droplet loading device are placed between top lid and bottom chamber. (b) Top view of the droplet loading device. The microchannel consists of a series of loading channels with a width of 200 µm. (c) During droplet loading (left), the loading channels are aligned above the microwells. After the droplet loading (right), the loading device is relocated to cover the microwells.

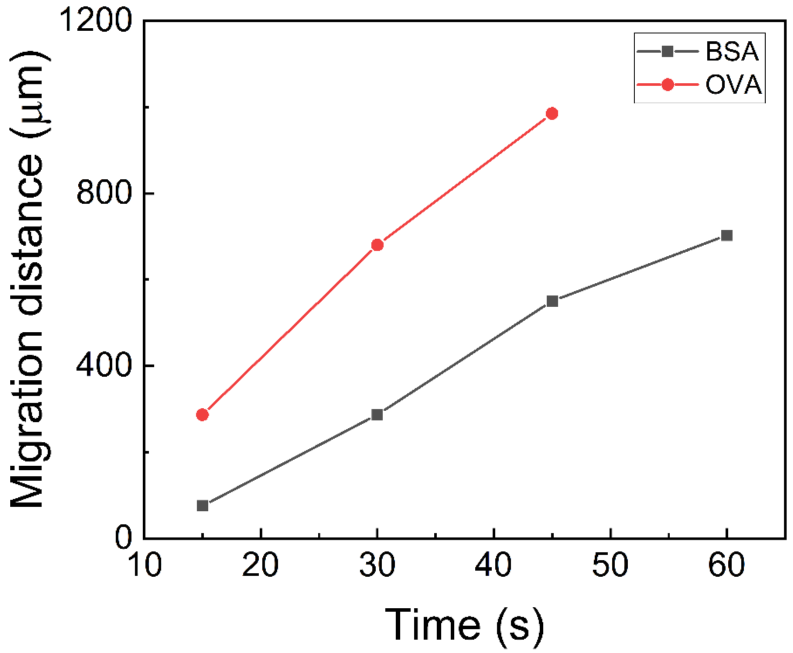


**Figure S3.** Electromigration of BSA and OVA over time (experiment). E: 40 V/cm;  Microwell diameter: 50 µm; Droplet diameter: 45 µm.

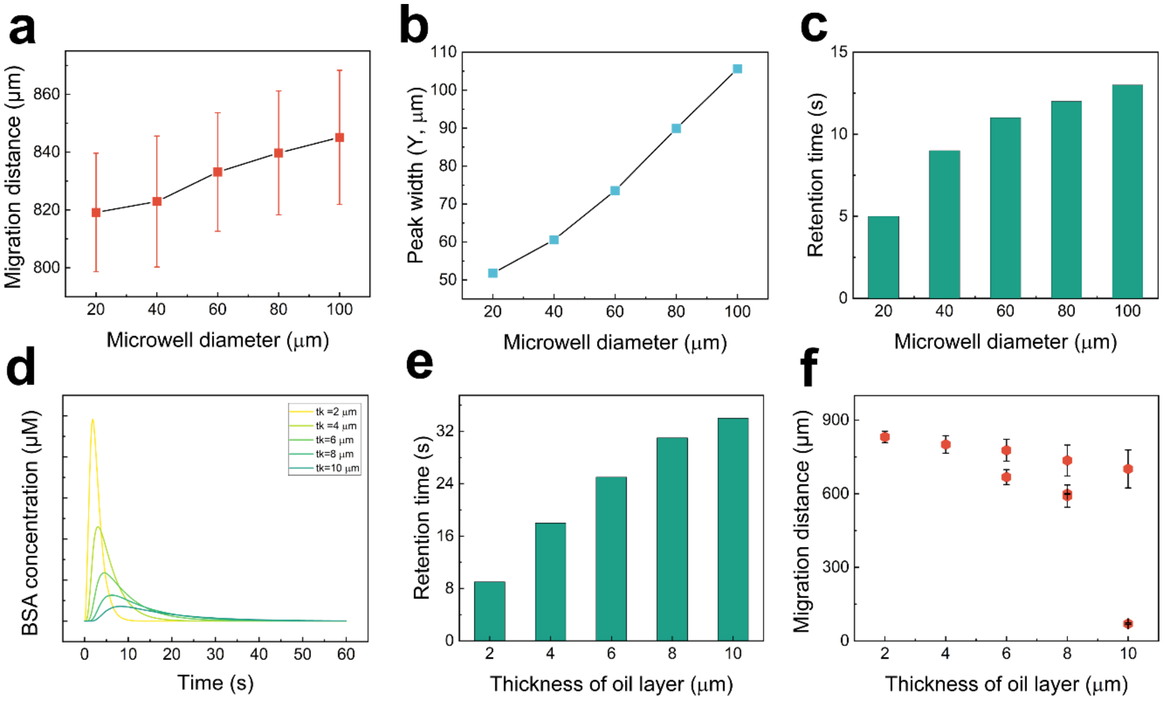


**Figure S4.** Simulation of the peak center/migration distance (a), peak width (b), and retention time (c, the time all proteins migrate out of microwells) of BSA with different microwell diameters (20-100 μm). The droplet was placed at the center of the microwell, and the oil layer thickness was set to 2.5 μm. the oil layer thickness is defined as the distance between the right edge of droplet and the left edge of microwell. Simulation of the effect of oil thickness on the change of BSA concentration at the right edge of microwell (d), retention time (e), and peak center/migration distance of BSA (f).

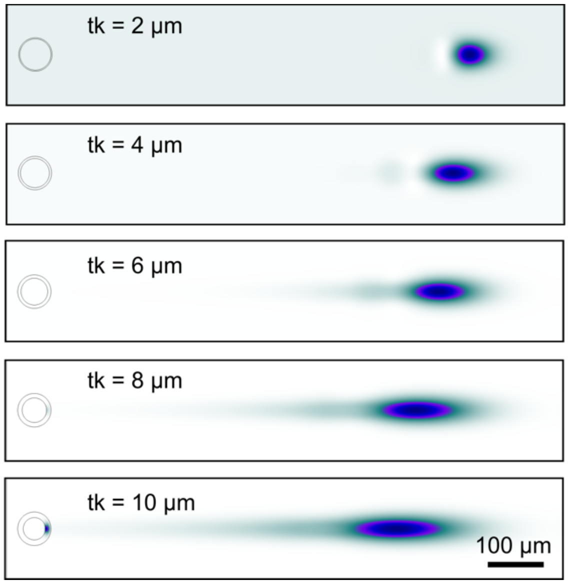


**Figure S5.** Electromigration of BSA with different thicknesses (tk) of oil layer (simulation). The microwell diameter is 60 μm. The electric field strength is 40 V/cm. t = 60s.

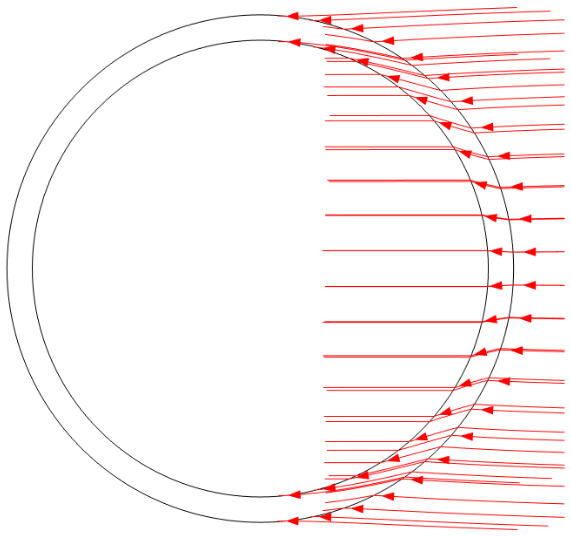

**Figure S6.** Streamline plot of current density near the interface between droplet, oil, and PA-gel (simulation). The droplet (inner circle) is positioned at the center of the microwell (outer circle). Electric field strength is 40 V/cm. The arrow represents current direction.

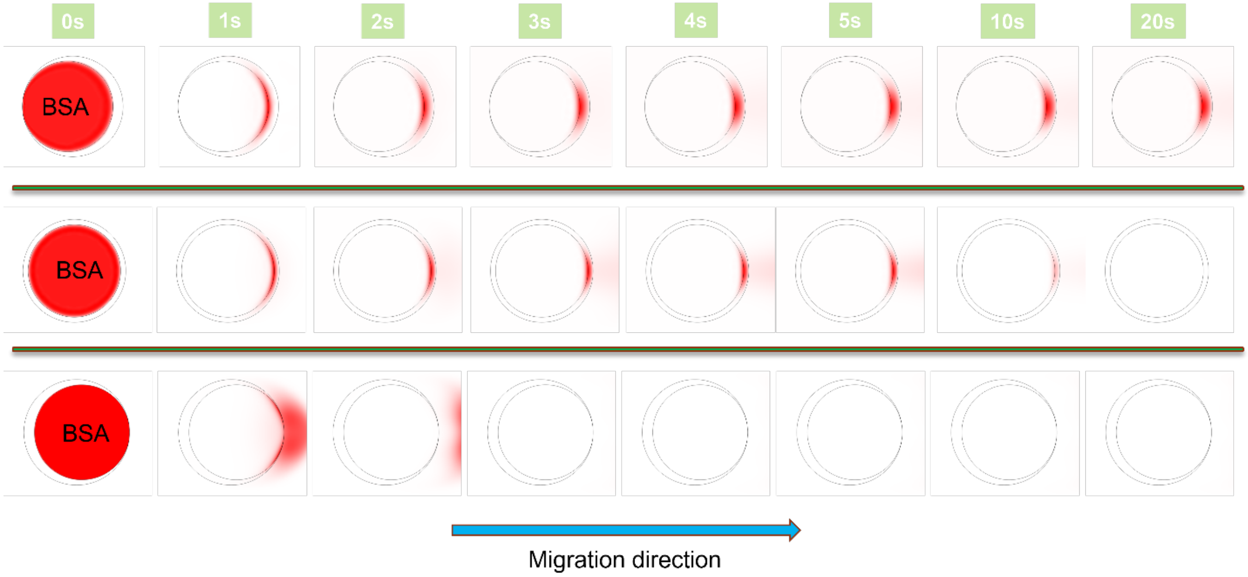


**Figure S7.** Electromigration of BSA with different relative positions of droplets (simulation). From the top to the bottom panel, the relative position of the droplet is right, middle, and right to the microwell, respectively. The electric field strength is 40 V/cm and the electrophoresis time is 0-20s.

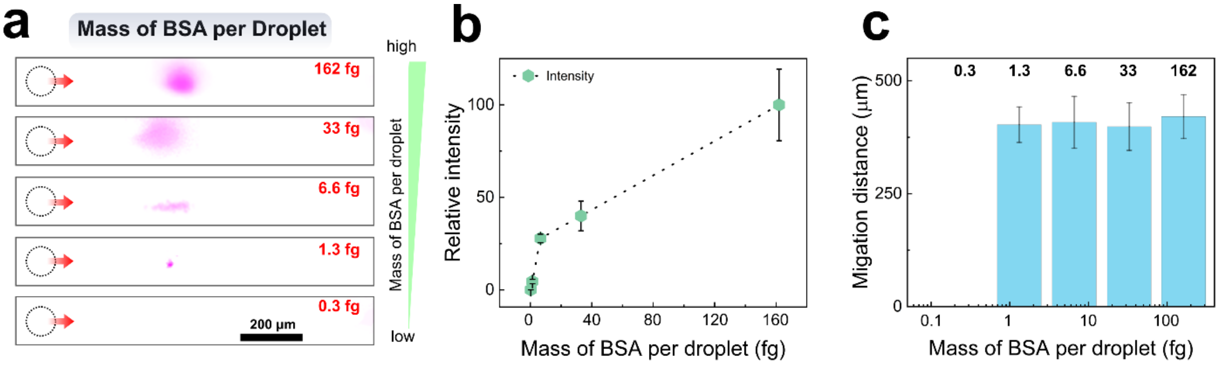


**Figure S8.** Limit of detection (LOD) of DropBlot on BSA analysis. (a) Immunofluorescence images, (b) Relative intensity, and (c) migration distance of BSA after 30s' electrophoresis with different protein mass per droplet (162 fg, 33 fg, 6.6fg, 1.3 fg, and 0.3 fg). Droplet diameter: 45 µm.

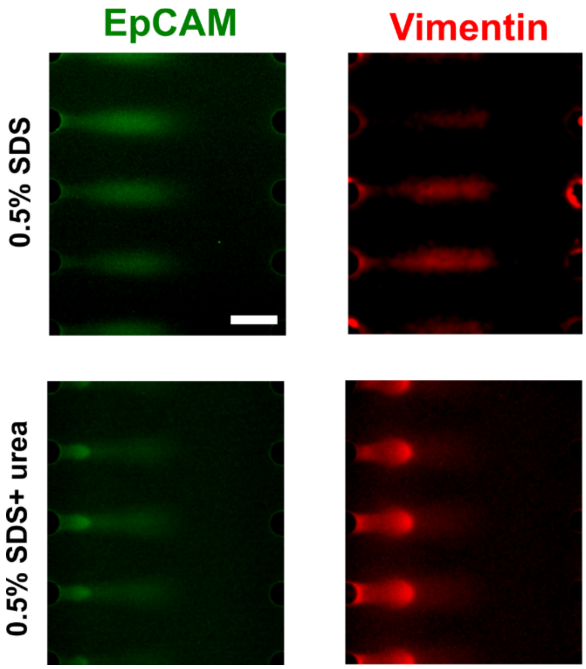


**Figure S9.** Intensity profiles of EpCAM (Green, MCF7) and Vimentin (Red, MDA-MB-231) when using an antigen-retrieval buffer containing 0.5% SDS only (top panel) and 0.5% SDS+ 6M urea (bottom panel), both after 30s electrophoresis at an electric field strength of 40V/cm. Droplet diameter: 45 μm. Scale bar: 200 μm.

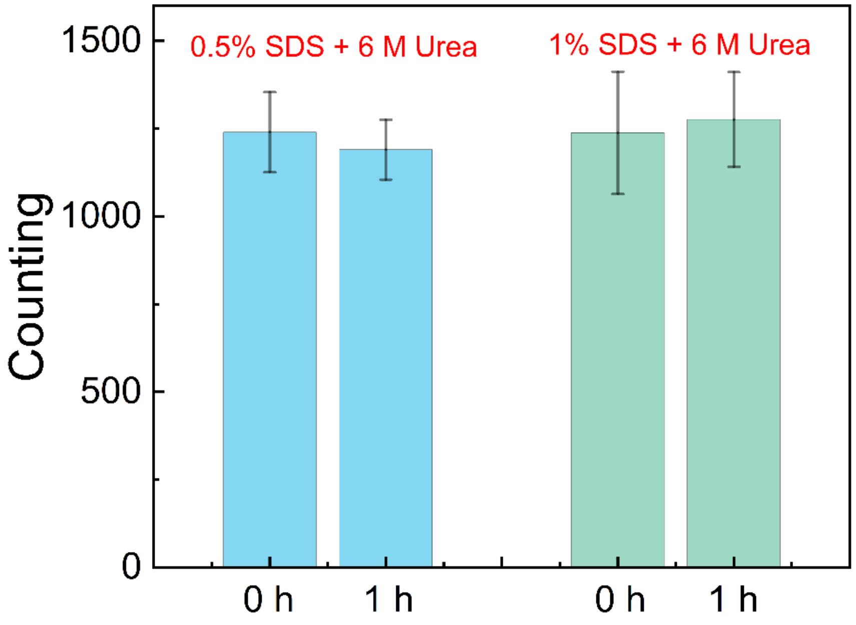


**Figure S10**. Droplet enumeration after 1-hour incubations at 100ºC. Droplets are loaded with 0.5% (w/v) SDS & 6 M Urea or 1% (w/v) SDS & 6 M Urea. Droplet diameter: 45 μm.

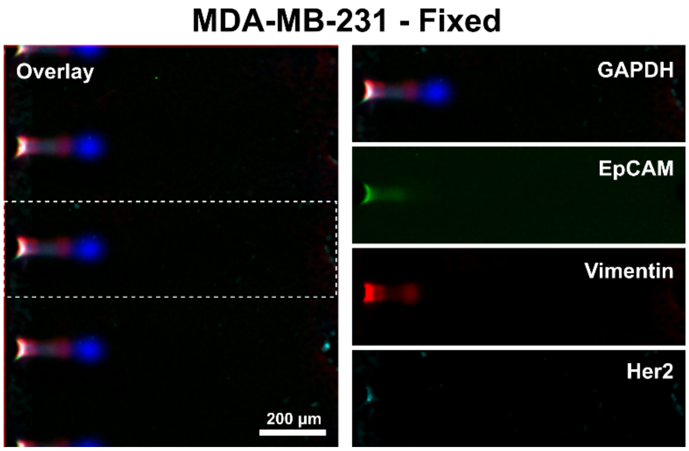


**Figure S11.** Immunofluorescence image of PFA-fixed MDA-MB-231 cells. Cells were fixed with 4% PFA at room temperature for 30min. Electric field strength: 60V/cm. Electrophoresis time: 30s.

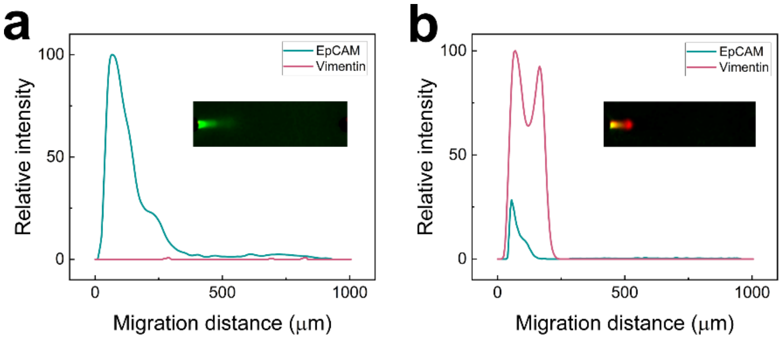


**Figure S12.** Immunofluorescence image and intensity profile of EpCAM in methanol-fixed MCF7 (a) and vimentin in methanol-fixed MDA-MB-231(b). Electric field strength: 60V/cm. Electrophoresis time: 60s.

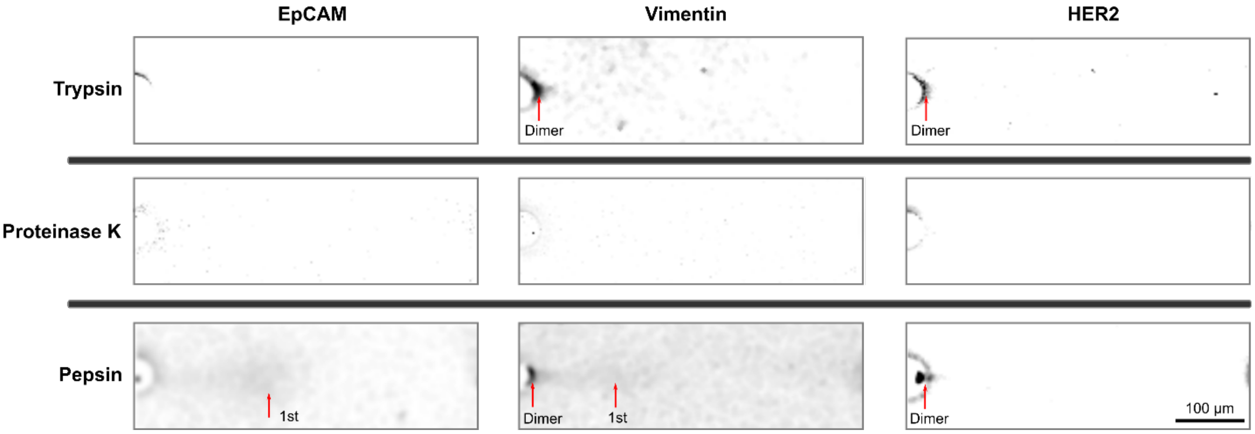


**Figure S13.** Enzymatic antigen retrieval from PFA-fixed MCF7 using trypsin, proteinase K, and pepsin. The cells were fixed with 4% PFA for 15 min at room temperature. Electric field strength: 60V/cm. Electrophoresis time: 30s.

### Table S1: Current Single-Cell Proteomics Analysis Techniques

| Name | Method | Fixed/ Live | Target | Surface /Intracellular Protein | Proteoforms detection | Throughput | Multiplexity | Pros | Cons | Ref |
|---|---|---|---|---|---|---|---|---|---|---|
| **Mass Spectrometry (Top down)** | Direct analysis of intact proteins | Live (fixed cells need new lysis step) | Protein | Both | Yes | <100 cells | >1000 | able to intact protein molecules, high sensitivity. | Low throughput, Limited to small-to-intermediate proteins (<25 kDa); difficulty to distinguish proteoforms due to high sample complexity | 1-3 |
| **Mass Spectrometry (Bottom up)** | Direct analysis of digested proteins | Live (fixed cells need new lysis step) | Protein | Both | Yes | <100 cells | >1000 | Highly multiplexed | Low throughput, limited to high abundant proteins (>10,000 copies /cell); Protein digestions will miss proteoform stoichiometry | 4-6 |
| **cyTOF (Cytometry by time of flight)** | Flow cytometry and inductive coupled plasma mass spectrometry; Use metal isotope tagged antibodies. | fixed cells, fixed tissues | Protein | both, intracellular proteins analysis requires cell fixation | Yes, based on the availability of proteoform antibody | ~ 100 cells /s | ~100 | Highly multiplexed (>100 protein targets); Low background | Cannot be applied to live cells; Limited availability of commercial metal-isotope-labelled antibodies; difficult to analyze single cells due to low recovery rate and high sample loss. | 7-8 |
| **FCM (flow cytometry)** | Label based, rely on fluorescent signals | live or fixed | Protein and nucleic acids | Both | Yes, based on the availability of proteoform antibody | 100-10,000 cells /s | ~17 | high throughput, highly multiplexed | Requires large sample. Low sensitivity due to spectra overlap and autofluorescence. Limited proteoform antibodies | 9-11 |
| **IMC (Imaging Mass Cytometry** | Labeled with isotope conjugated antibodies, and analyzed with Mass Spectrometry) | Fixed / Frozen Tissue | Protein | Both | Yes | 1 mm^2 / 2h | ~40 | High sensitivity, highly multiplexed; advance in spatial resolution | Low throughput; Limited to small proteins (<20 kDa), Limited proteoform antibodies; High Cost; | 12-13 |
| **(PiMS) Proteoform Imaging Mass Spectrometry** | Combination of nanospray desorption electrospray (nano-DESI) and individual ion MS (I^2 MS) | Fixed Tissue | Protein | Both | Yes | 2.5-4 um /s | ~169 (proteoforms) | highly multiplexed; High spatial resolution | Limited to proteins smaller than 70 kDa | 14 |
| **Abseq** | label-based, antibodies are labeled with sequence tags | Live (compatible to fixed cellss) | Protein | Surface | Not applied yet, but in theory it can. | 10000 cells / 1h | unlimited, but can be reduced based on the available proteins, and reading capacity) | high sensitivity to low-abundance antibodies (single molecule per cell); | some antibodies cannot be labeled with detectable tags | 15 |
| **On-Chip Cytometry (Microengraving)** | Label-Based, rely on fluorescent signals. | Live | Protein and nucleic acids | Surface | Yes | 84,672 cells/ array | ~4 | parallel study; Cells can be recovered. | Limited to availability of proteoform antibodies. | 16 |
| **Single-cell barcode chips (SCBCs)** | Label-Based, rely on fluorescent signals. | Live/Fix | Protein | secreted proteins | Not applied yet, but in theory it can. | 3000-5000 cells/ array | ~42 | parallel study; Cells can be recovered. Highly multiplexed | Limited to secreted proteins | 17 |
| **Quantitive Ferrohydrodynamic Cell Separation (qFCS)** | Label-based, cells are labeld with magnetic beads and sorted based on the antigen density | Live/Fix | Protein | Surface | No | 30,000 cells/min | 1 | can detect rare cell types, as low as 10 cells/mL; high sensitivity to low abundance antigens. | Low multiplexed. Cannot analyze proteoforms. | 18 |

| | | | | | | | | | | |
|---|---|---|---|---|---|---|---|---|---|---|
| **single cell western blot (scWB, 2D)** | First separate based on molecular weight, and then use fluorescent antibodies to visualize protein targets | Live | Protein | Both | Yes | ~5000 cells/ array | ~12 | Parallel study; Capable of analyzing proteoforms. | Low multiplexed. | 19-20 |
| **single cell western blot (scWB, 3D)** | First separate based on molecular weight, and then use fluorescent antibodies to visualize protein targets | Live | Protein | Both | Yes | 2.5 cells /s, 300 cells/ array | ~4 | Parallel study; Low sample consumption | Low multiplexed. A large number of images to process. Cannot be applied to fixed cells | 21 |
| **Magnetic ranking cytometry (MagRC)** | Label-based, cells are labeled with magnetic nanoparticles. | Live/Fix | Protein | Surface | No | 500 ul/h | 1 | suitable to rare cells. | Low multiplexed. | 22 |
| **Droplet-based cell screening & sorting** | Label-based, proteins are labeled with fluorescence antibodies | Live | Protein | Surface | No | 2 - 5e5 cells/ h | 1 | high sensitivity, minimal cross-contamination | Low multiplexed. | 23-24 |
| **Digital microfluidics (DMF)** | Digital microfluidics provide single cell sample (protein, nucleic acids) for downstream analysis (e.g., LC-MS/MS); Sample preparation is in droplet | Live/Fix | Protein/nucleic acids | Both | Not applied yet, but in theory it can. | 50-500 cell / assay | >1000 | High precision, low sample consumption and ability of perform complex manipulation of small volumes of liquid | Low throughput; complicated; target detection relies on other techniques (e.g., MS). | 25-27 |
| **oil-air droplet (OAD) chip** | Combination of droplet microfluidics and LC-MS/MS | Live | Protein | Both | Not applied yet, but in theory it can. | 1-100 cells/ chip | ~355 | Low sample loss, high sample injection efficiency | Low throughput; complicated; target detection relies on other techniques (e.g, MS). | 28 |
| **Nanodroplet processing in one-pot (nanoPOTS)** | Combination of droplet microfluidics and LC-MS/MS | Live | Protein | Both | Not applied yet, but in theory it can. | 10-240 cells / assay | 670-3000 | highly multiplexed; low sample contamination | Low throughput; complicated; target detection relies on other techniques (e.g., MS). | 6, 29 |
| **Single-cell integrated proteomic microfluidic chip (SciProChip)** | Combination of on-chip peptide preparation and LC-MS/MS | Live | Protein | Both | Not applied yet, but in theory it can. | 20 / assay | ~1500 | highly multiplexed; low sample contamination | Low throughput; complicated; target detection relies on other techniques (e.g., MS). | 30 |
| **Immunohistochemistry (IHC) or Immunocytochemistry (ICC)** | Label-based, proteins are labeled with antibodies and visualized with colored chromogen or fluorophores | Fixed | Proteins | Both | Yes | NA | ~2-50 | High specificity, tissue localization, wide applicability, | Limited by the availability of antibodies, narrow dynamic range, time consuming, variability | 31-33 |

### Table S2. Proteins analyzed in DropBlot

| Name | MW (kDa) | Function (normal) | Locations | Isoforms | Isoform MW (kDa) | Isoform Formation & Function |
|---|---|---|---|---|---|---|
| **GAPDH** | 36 | Glycolytic enzyme, regulate mRNA stability | Mainly in cytoplasm, also founded in nucleus, mitochondria, cell membrane[34-35] | Yes, NA | NA | Not studied here |
| **Vimentin** | 55 | Structural protein, maintain cellular integrity and provide resistance against stress, cellular signal transduction[36] | Mainly in cytoplasm, can b on the surface of activated platelets (secreted by activated macrophage)[37], also found on the plasma membrane[38] | #1[39] | 150, dimer | the 150 kDa band likely represents two covalently linked monomers originally belonging to adjacent dimers or tetramers that dissociate in SDS-PAGE |
| | | | | #2[40] | 120 | a high molecular weight form of vimentin, 120 kDa, within and adjacent to vesicles near the luminal surface of Human Microvascular Endothelial Cells (HMEC) |
| | | | | #3[38] | 60 | membrane-associated 60 kDa vimentin isoform is also present in membranes of non-activated lymphocytes, which cannot bind extracellular anti-vimentin antibodies |
| | | | | #4[41] | 49 | activated human T cells |
| | | | | #5[42] | 47 | Spliced variant, ~35 amino acids smaller. |
| | | | | #6[43] | 32 | Fragmentation pattern of vimentin due to cell apoptosis |
| | | | | #7[43] | 20 | Fragmentation pattern of vimentin due to cell apoptosis |
| **EpCAM** | 40 | Cell adhesion protein, cell signaling, proliferation, differentiation[44] | Cell membrane | #1[45] | 66, dimer | extracellular part of human EpCAM forms a heart-shaped dimer, which would form at cell surfaces |
| | | | | #2[46-47] | 35 | epithelium-like tumor cell lines MCF-7, T47D, and SkBR3 showed strong expression of the EpCAM protein as basic and glycosylated isoforms of 35 |
| | | | | #3[48] | 32 | Proteolytic cleavage |
| | | | | #4[48] | 6 | Proteolytic cleavage |
| **Her2** | 185 | Provides the cell with potent proliferative and anti-apoptosis signals[49] | Cell membrane | Yes, NA | - | Not studied here |

**Table S3. Formalin vs. Methanol Fixation**

| Fixation Method | Formalin/PFA | Methanol |
|---|---|---|
| **Composition of Fixative** | Formaldehyde solution in water | Methanol |
| **Chemical Reaction** | Crosslinks proteins with adjacent amino acids, nucleic acids, and lipids | Lipids are removed from membranes, proteins precipitate. |
| **Penetration** | Slower | Faster |
| **Cell Morphology** | Preserve morphology well | Can cause tissue shrinkage and distortion |
| **Antigenicity** | May alter or mask some antigens | Generally preserve antigenicity |
| **Enzymatic Activity** | May preserve some enzymatic activity | Generally preserve enzymatic activity |
| **Nucleic Acids** | May alter or mask nucleic acids | Generally preserve nucleic acids |
| **Other concerns** | Protein can move through the cell during fixation, so you can see nuclear protein in cytosol | Small soluble metabolites leak. |
| **Applications** | Histology, immunohistochemistry, in situ hybridization, suitable to tissue | Cytology, immunofluorescence, flow cytometry, suitable to cell |

### Table S4. Antigen Retrieval from 4% PFA-fixed MCF7 using Enzymes

| **Enzyme** | **Trypsin** | **Trypsin** | **Trypsin** | **Trypsin** | **Trypsin** | **Trypsin** | **Proteinase K** | **Proteinase K** | **Pepsin** | **Pepsin** | **Pepsin** |
|---|---|---|---|---|---|---|---|---|---|---|---|
| **Incubation condition** | fixed cells, lysis buffe, and Trypsin were encapsulated In droplet, incubate for 30 at RT | fixed cells are incubated with trypsin solution in 1.5 mL Eppendorf tube, 30 min at RT. After incubation, wash with PBS for 3 times. | fixed cells are incubated with trypsin solution in 1.5 mL Eppendorf tube, 15 min at RT. After incubation, wash with PBS for 3 times. | fixed cells are incubated with trypsin solution in 1.5 mL Eppendorf tube, 15 min at RT. After incubation, wash with PBS for 3 times. | fixed cells are incubated with trypsin solution in 1.5 mL Eppendorf tube, 10 min at RT. After incubation, wash with PBS for 3 times. | fixed cells are incubated with trypsin solution in 1.5 mL Eppendorf tube, 5 min at RT. After incubation, wash with PBS for 3 times. | fixed cells are incubated with trypsin solution in 1.5 mL Eppendorf tube, 7 min at RT. After incubation, wash with PBS for 3 times. | fixed cells are incubated with trypsin solution in 1.5 mL Eppendorf tube, 7 min at RT. After incubation, wash with PBS for 3 times. | fixed cells are incubated with trypsin solution in 1.5 mL Eppendorf tube, 10 min at 37. After incubation, wash with PBS for 3 times. | fixed cells are incubated with trypsin solution in 1.5 mL Eppendorf tube, 10 min at 37. After incubation, wash with PBS for 3 times. | fixed cells are incubated with trypsin solution in 1.5 mL Eppendorf tube, 15 min at 37. After incubation, wash with PBS for 3 times. |
| **Lysis condition** | 0.5%(w/v) SDS + 6M Urea, 30 min at RT | 0.5%(w/v) SDS + 6M Urea, 30 min at RT | 0.5%(w/v) SDS + 6M Urea, 30 min at RT | 0.5%(w/v) SDS + 6M Urea, 30 min at 98C | 0.5%(w/v) SDS + 6M Urea, 30 min at RT | 0.5%(w/v) SDS + 6M Urea, 30 min at RT | 0.5%(w/v) SDS + 6M Urea, 30 min at RT | 0.5%(w/v) SDS + 6M Urea, 30 min at 98C | 0.5%(w/v) SDS + 6M Urea, 30 min at RT | 0.5%(w/v) SDS + 6M Urea, 30 min at 98C | 0.5%(w/v) SDS + 6M Urea, 30 min at RT |
| **EpCAM (Y/N)** | N | N | N | N | N | N | N | N | N | N | **weak , no isoform** |
| **Vimentin (Y/N)** | N | N | N | Y (Dimer) | N | N | N | N | Y | N | **Y** |
| **Her2 (Y/N)** | N | N | N | Y (Dimer) | N | N | N | N | Y (Dimer) | N | **Y (Dimer)** |

**Table S5. Samples Tested with DropBlot**

| Patient | ID | ER-α Status | PR, HER2 Status | Cell Status | Type | Signal |
|---|---|---|---|---|---|---|
| **1** | 041318 | ER-$\alpha^{3+}$ | PR+, $HER2^{-}$ | Suspension, Fresh | Invasive ductal breast tumor, PT2pN0 | Yes |
| **2** | 121715 | ER-$\alpha^{1+}$ | PR-, $HER2^{+}$ | Suspension, Fresh | Lymph node infiltrated breast tumor | Yes |
| **3** | 041318 | ER-$\alpha^{3+}$ | PR+, $HER2^{-}$ | Tissue, Fresh | Invasive ductal breast tumor | Yes |
| **4** | 102816 | ER-$\alpha^{1+}$ | PR+, $HER2^{+}$ | Tissue, Fresh | Triple positive breast tumor | Yes |
| **5** | 040615 | - | - | Tissue, Fresh | Cureline Bca Fresh Tissue | Yes |
| **6** | 32818-5 | ER-$\alpha^{1+}$ | PR+, $HER2^{-}$ | Suspension, Fresh | Invasive ductal breast tumor | No |
| **7** | 121615 | ER-$\alpha^{3+}$ | $PR^{3+}$, $HER2^{-}$ | Suspension, Fresh | Lymph node infiltrated breast tumor, T4bN1A | No |
| **8** | 032918 | ER-α- | PR+, $HER2^{-}$ | Suspension, Fresh | Breast tumor | No |
| **9** | 062615 | - | $HER2^{+}$ | FFPE | Breast tumor | No |
| **10** | 062615 | - | $HER2^{+}$ | FFPE | Breast tumor | No |
| **11** | 062615 | - | $HER2^{+}$ | FFPE | Breast tumor | No |

### References


1. van Remoortere, A.; van Zeijl, R. J.; van den Oever, N.; Franck, J.; Longuespee, R.; Wisztorski, M.; Salzet, M.; Deelder, A. M.; Fournier, I.; McDonnell, L. A., Maldi Imaging and Profiling Ms of Higher Mass Proteins from Tissue. *J Am Soc Mass Spectrom* **2010,** *21* (11), 1922-9.
2. Aichler, M.; Walch, A., Maldi Imaging Mass Spectrometry: Current Frontiers and Perspectives in Pathology Research and Practice. *Lab Invest* **2015,** *95* (4), 422-31.
3. Do, T. D.; Ellis, J. F.; Neumann, E. K.; Comi, T. J.; Tillmaand, E. G.; Lenhart, A. E.; Rubakhin, S. S.; Sweedler, J. V., Optically Guided Single Cell Mass Spectrometry of Rat Dorsal Root Ganglia to Profile Lipids, Peptides and Proteins. *Chemphyschem* **2018,** *19* (10), 1180-1191.
4. Budnik, B.; Levy, E.; Harmange, G.; Slavov, N., Scope-Ms: Mass Spectrometry of Single Mammalian Cells Quantifies Proteome Heterogeneity During Cell Differentiation. *Genome Biol* **2018,** *19* (1), 161.
5. Zhu, Y.; Clair, G.; Chrisler, W. B.; Shen, Y.; Zhao, R.; Shukla, A. K.; Moore, R. J.; Misra, R. S.; Pryhuber, G. S.; Smith, R. D.; Ansong, C.; Kelly, R. T., Proteomic Analysis of Single Mammalian Cells Enabled by Microfluidic Nanodroplet Sample Preparation and Ultrasensitive Nanolc-Ms. *Angew Chem Int Ed Engl* **2018,** *57* (38), 12370-12374.
6. Zhu, Y.; Piehowski, P. D.; Zhao, R.; Chen, J.; Shen, Y.; Moore, R. J.; Shukla, A. K.; Petyuk, V. A.; Campbell-Thompson, M.; Mathews, C. E.; Smith, R. D.; Qian, W. J.; Kelly, R. T., Nanodroplet Processing Platform for Deep and Quantitative Proteome Profiling of 10-100 Mammalian Cells. *Nat Commun* **2018,** *9* (1), 882.
7. Angelo, M.; Bendall, S. C.; Finck, R.; Hale, M. B.; Hitzman, C.; Borowsky, A. D.; Levenson, R. M.; Lowe, J. B.; Liu, S. D.; Zhao, S.; Natkunam, Y.; Nolan, G. P., Multiplexed Ion Beam Imaging of Human Breast Tumors. *Nat Med* **2014,** *20* (4), 436-42.
8. Bandura, D. R.; Baranov, V. I.; Ornatsky, O. I.; Antonov, A.; Kinach, R.; Lou, X.; Pavlov, S.; Vorobiev, S.; Dick, J. E.; Tanner, S. D., Mass Cytometry: Technique for Real Time Single Cell Multitarget Immunoassay Based on Inductively Coupled Plasma Time-of-Flight Mass Spectrometry. *Anal Chem* **2009,** *81* (16), 6813-22.
9. Irish, J. M.; Myklebust, J. H.; Alizadeh, A. A.; Houot, R.; Sharman, J. P.; Czerwinski, D. K.; Nolan, G. P.; Levy, R., B-Cell Signaling Networks Reveal a Negative Prognostic Human Lymphoma Cell Subset That Emerges During Tumor Progression. *P Natl Acad Sci USA* **2010,** *107* (29), 12747-12754.
10. Nuti, E.; Rossello, A.; Cuffaro, D.; Camodeca, C.; Van Bael, J.; van der Maat, D.; Martens, E.; Fiten, P.; Pereira, R. V. S.; Ugarte-Berzal, E.; Gouwy, M.; Opdenakker, G.; Vandooren, J., Bivalent Inhibitor with Selectivity for Trimeric Mmp-9 Amplifies Neutrophil Chemotaxis and Enables Functional Studies on Mmp-9 Proteoforms. *Cells-Basel* **2020,** *9* (7).
11. Krutzik, P. O.; Crane, J. M.; Clutter, M. R.; Nolan, G. P., High-Content Single-Cell Drug Screening with Phosphospecific Flow Cytometry. *Nat Chem Biol* **2008,** *4* (2), 132-142.
12. Jackson, H. W.; Fischer, J. R.; Zanotelli, V. R. T.; Ali, H. R.; Mechera, R.; Soysal, S. D.; Moch, H.; Muenst, S.; Varga, Z.; Weber, W. P.; Bodenmiller, B., The Single-Cell Pathology Landscape of Breast Cancer. *Nature* **2020,** *578* (7796), 615-+.
13. Baharlou, H.; Canete, N. P.; Cunningham, A. L.; Harman, A. N.; Patrick, E., Mass Cytometry Imaging for the Study of Human Diseases-Applications and Data Analysis Strategies. *Front Immunol* **2019,** *10*.
14. Su, P.; McGee, J. P.; Durbin, K. R.; Hollas, M. A. R.; Yang, M.; Neumann, E. K.; Allen, J. L.; Drown, B. S.; Butun, F. A.; Greer, J. B.; Early, B. P.; Fellers, R. T.; Spraggins, J. M.; Laskin, J.; Camarillo, J. M.;

Kafader, J. O.; Kelleher, N. L., Highly Multiplexed, Label-Free Proteoform Imaging of Tissues by Individual Ion Mass Spectrometry. *Sci Adv* **2022,** *8* (32), eabp9929.
15. Shahi, P.; Kim, S. C.; Haliburton, J. R.; Gartner, Z. J.; Abate, A. R., Abseq: Ultrahigh-Throughput Single Cell Protein Profiling with Droplet Microfluidic Barcoding. *Sci Rep* **2017,** *7*, 44447.
16. Ogunniyi, A. O.; Thomas, B. A.; Politano, T. J.; Varadarajan, N.; Landais, E.; Poignard, P.; Walker, B. D.; Kwon, D. S.; Love, J. C., Profiling Human Antibody Responses by Integrated Single-Cell Analysis. *Vaccine* **2014,** *32* (24), 2866-73.
17. Lu, Y.; Xue, Q.; Eisele, M. R.; Sulistijo, E. S.; Brower, K.; Han, L.; Amir el, A. D.; Pe'er, D.; Miller-Jensen, K.; Fan, R., Highly Multiplexed Profiling of Single-Cell Effector Functions Reveals Deep Functional Heterogeneity in Response to Pathogenic Ligands. *Proc Natl Acad Sci U S A* **2015,** *112* (7), E607-15.
18. Liu, Y.; Vieira, R. M. S.; Mao, L., Simultaneous and Multimodal Antigen-Binding Profiling and Isolation of Rare Cells Via Quantitative Ferrohydrodynamic Cell Separation. *ACS Nano* **2023,** *17* (1), 94-110.
19. Hughes, A. J.; Spelke, D. P.; Xu, Z.; Kang, C. C.; Schaffer, D. V.; Herr, A. E., Single-Cell Western Blotting. *Nat Methods* **2014,** *11* (7), 749-55.
20. Sinkala, E.; Sollier-Christen, E.; Renier, C.; Rosas-Canyelles, E.; Che, J.; Heirich, K.; Duncombe, T. A.; Vlassakis, J.; Yamauchi, K. A.; Huang, H.; Jeffrey, S. S.; Herr, A. E., Profiling Protein Expression in Circulating Tumour Cells Using Microfluidic Western Blotting. *Nat Commun* **2017,** *8*, 14622.
21. Grist, S. M.; Mourdoukoutas, A. P.; Herr, A. E., 3d Projection Electrophoresis for Single-Cell Immunoblotting. *Nat Commun* **2020,** *11* (1), 6237.
22. Poudineh, M.; Aldridge, P. M.; Ahmed, S.; Green, B. J.; Kermanshah, L.; Nguyen, V.; Tu, C.; Mohamadi, R. M.; Nam, R. K.; Hansen, A.; Sridhar, S. S.; Finelli, A.; Fleshner, N. E.; Joshua, A. M.; Sargent, E. H.; Kelley, S. O., Tracking the Dynamics of Circulating Tumour Cell Phenotypes Using Nanoparticle-Mediated Magnetic Ranking. *Nat Nanotechnol* **2017,** *12* (3), 274-281.
23. Mazutis, L.; Gilbert, J.; Ung, W. L.; Weitz, D. A.; Griffiths, A. D.; Heyman, J. A., Single-Cell Analysis and Sorting Using Droplet-Based Microfluidics. *Nat Protoc* **2013,** *8* (5), 870-91.
24. Martino, C.; Zagnoni, M.; Sandison, M. E.; Chanasakulniyom, M.; Pitt, A. R.; Cooper, J. M., Intracellular Protein Determination Using Droplet-Based Immunoassays. *Anal Chem* **2011,** *83* (13), 5361-8.
25. Eyer, K.; Doineau, R. C. L.; Castrillon, C. E.; Briseno-Roa, L.; Menrath, V.; Mottet, G.; England, P.; Godina, A.; Brient-Litzler, E.; Nizak, C.; Jensen, A.; Griffiths, A. D.; Bibette, J.; Bruhns, P.; Baudry, J., Single-Cell Deep Phenotyping of Igg-Secreting Cells for High-Resolution Immune Monitoring. *Nat Biotechnol* **2017,** *35* (10), 977-982.
26. Steinbach, M. K.; Leipert, J.; Blurton, C.; Leippe, M.; Tholey, A., Digital Microfluidics Supported Microproteomics for Quantitative Proteome Analysis of Single Caenorhabditis Elegans Nematodes. *J Proteome Res* **2022,** *21* (8), 1986-1996.
27. Leipert, J.; Steinbach, M. K.; Tholey, A., Isobaric Peptide Labeling on Digital Microfluidics for Quantitative Low Cell Number Proteomics. *Anal Chem* **2021,** *93* (15), 6278-6286.
28. Li, Z. Y.; Huang, M.; Wang, X. K.; Zhu, Y.; Li, J. S.; Wong, C. C. L.; Fang, Q., Nanoliter-Scale Oil-Air-Droplet Chip-Based Single Cell Proteomic Analysis. *Anal Chem* **2018,** *90* (8), 5430-5438.
29. Cong, Y.; Motamedchaboki, K.; Misal, S. A.; Liang, Y.; Guise, A. J.; Truong, T.; Huguet, R.; Plowey, E. D.; Zhu, Y.; Lopez-Ferrer, D.; Kelly, R. T., Ultrasensitive Single-Cell Proteomics Workflow Identifies >1000 Protein Groups Per Mammalian Cell. *Chem Sci* **2020,** *12* (3), 1001-1006.
30. Gebreyesus, S. T.; Siyal, A. A.; Kitata, R. B.; Chen, E. S.; Enkhbayar, B.; Angata, T.; Lin, K. I.; Chen, Y. J.; Tu, H. L., Streamlined Single-Cell Proteomics by an Integrated Microfluidic Chip and Data-Independent Acquisition Mass Spectrometry. *Nat Commun* **2022,** *13* (1), 37.
31. Ramos-Vara, J. A., Technical Aspects of Immunohistochemistry. *Vet Pathol* **2005,** *42* (4), 405-26.
32. Happonen, R. P.; Heikinheimo, K., Introduction to Immunocytochemistry. *Proc Finn Dent Soc* **1989,** *85* (2), 61-7.

33. Hofman, P.; Badoual, C.; Henderson, F.; Berland, L.; Hamila, M.; Long-Mira, E.; Lassalle, S.; Roussel, H.; Hofman, V.; Tartour, E.; Ilie, M., Multiplexed Immunohistochemistry for Molecular and Immune Profiling in Lung Cancer-Just About Ready for Prime-Time? *Cancers (Basel)* **2019,** *11* (3).

34. Nicholls, C.; Li, H.; Liu, J. P., Gapdh: A Common Enzyme with Uncommon Functions. *Clin Exp Pharmacol Physiol* **2012,** *39* (8), 674-9.

35. Tristan, C.; Shahani, N.; Sedlak, T. W.; Sawa, A., The Diverse Functions of Gapdh: Views from Different Subcellular Compartments. *Cell Signal* **2011,** *23* (2), 317-23.

36. Ivaska, J.; Pallari, H. M.; Nevo, J.; Eriksson, J. E., Novel Functions of Vimentin in Cell Adhesion, Migration, and Signaling. *Exp Cell Res* **2007,** *313* (10), 2050-62.

37. Mor-Vaknin, N.; Punturieri, A.; Sitwala, K.; Markovitz, D. M., Vimentin Is Secreted by Activated Macrophages. *Nat Cell Biol* **2003,** *5* (1), 59-63.

38. Bilalic, S.; Michlmayr, A.; Gruber, V.; Buchberger, E.; Burghuber, C.; Bohmig, G. A.; Oehler, R., Lymphocyte Activation Induces Cell Surface Expression of an Immunogenic Vimentin Isoform. *Transpl Immunol* **2012,** *27* (2-3), 101-106.

39. Monico, A.; Guzman-Caldentey, J.; Pajares, M. A.; Martin-Santamaria, S.; Perez-Sala, D., Molecular Insight into the Regulation of Vimentin by Cysteine Modifications and Zinc Binding. *Antioxidants-Basel* **2021,** *10* (7).

40. Xu, B.; deWaal, R. M.; Mor-Vaknin, N.; Hibbard, C.; Markovitz, D. M.; Kahn, M. L., The Endothelial Cell-Specific Antibody Pal-E Identifies a Secreted Form of Vimentin in the Blood Vasculature. *Mol Cell Biol* **2004,** *24* (20), 9198-9206.

41. Rose, M. L., Role of Anti-Vimentin Antibodies in Allograft Rejection. *Hum Immunol* **2013,** *74* (11), 1459-1462.

42. von Brandenstein, M.; Puetz, K.; Schlosser, M.; Loser, H.; Kallinowski, J. P.; Godde, D.; Buettner, R.; Storkel, S.; Fries, J. W. U., Vimentin 3, the New Hope, Differentiating Rcc Versus Oncocytoma. *Dis Markers* **2015,** *2015*.

43. Mary, S.; Kulkarni, M. J.; Mehendale, S. S.; Joshi, S. R.; Giri, A. P., Differential Accumulation of Vimentin Fragments in Preeclamptic Placenta. *Cytoskeleton* **2017,** *74* (11), 420-425.

44. Huang, L.; Yang, Y. H.; Yang, F.; Liu, S. M.; Zhu, Z. Q.; Lei, Z. L.; Guo, J., Functions of Epcam in Physiological Processes and Diseases (Review). *Int J Mol Med* **2018,** *42* (4), 1771-1785.

45. Pavsic, M.; Guncar, G.; Djinovic-Carugo, K.; Lenarcic, B., Crystal Structure and Its Bearing Towards an Understanding of Key Biological Functions of Epcam. *Nature Communications* **2014,** *5*.

46. Martowicz, A.; Spizzo, G.; Gastl, G.; Untergasser, G., Phenotype-Dependent Effects of Epcam Expression on Growth and Invasion of Human Breast Cancer Cell Lines. *Bmc Cancer* **2012,** *12*.

47. Schmidt, D. S.; Klingbeil, P.; Schnolzer, M.; Zoller, M., Cd44 Variant Isoforms Associate with Tetraspanins and Epcam. *Experimental Cell Research* **2004,** *297* (2), 329-347.

48. Blazar, M.; Briaire-De Bruijn, I. H.; Rees-Bakker, H. A. M.; Prins, F. A.; Helfrich, W.; de Leij, L.; Riethmuller, G.; Alberti, S.; Warnaar, S. O.; Fleuren, G. J.; Litvinov, S. V., Epidermal Growth Factor-Like Repeats Mediate Lateral and Reciprocal Interactions of Ep-Cam Molecules in Homophilic Adhesions. *Mol Cell Biol* **2001,** *21* (7), 2570-2580.

49. Gutierrez, C.; Schiff, R., Her2: Biology, Detection, and Clinical Implications. *Archives of pathology & laboratory medicine* **2011,** *135* (1), 55-62.
